# Integrated control of redox and energy metabolism by the membrane-bound and soluble transhydrogenases of *Pseudomonas putida* across metabolic regimes

**DOI:** 10.1101/2025.11.04.686620

**Authors:** Michele Partipilo, Giusi Favoino, Òscar Puiggené, Catarina Rocha, Carina Meiners, Nicolás Gurdo, Stefano Donati, Daniel C. Volke, Pablo I. Nikel

## Abstract

Redox homeostasis is central to microbial physiology and stress adaptation, yet the functional roles of transhydrogenases remain poorly understood beyond a few organisms. In this study, we systematically explored how *Pseudomonas putida*, a model soil bacterium, integrates two distinct transhydrogenases (membrane-bound PntAB and soluble SthA) into a flexible and reversible redox-balancing system that supports metabolic robustness across diverse metabolic regimes. While single deletions of either enzyme had minimal impact on the overall fitness, the double Δ*pntAB* Δ*sthA* mutant exhibited growth defects, disrupted energy charge, and redox imbalance. Unexpectedly, SthA proved essential for acetate-dependent growth, a phenotype traced to a transcriptional regulator involved in glyoxylate metabolism. Transhydrogenases also mediated tolerance to formate, a key one-carbon (C_1_) substrate for biotechnology, by channeling reducing equivalents released during feedstock oxidation. Synergistic activity with native formate dehydrogenases enabled redox buffering, even under stressful conditions. Functional complementation with native and engineered NAD^+^- or NADP^+^-dependent dehydrogenases validated SthA as the main sink for excess NADH. Comparative genomics linked transhydrogenase gene neighborhoods to stress and membrane processes, highlighting their evolutionary significance. These findings redefine transhydrogenases as dynamic regulators of redox metabolism, not passive cofactor shuttles. Furthermore, this work positions *P*. *putida* as a prime host for redox-intensive applications, informing design principles for C_1_-based metabolic engineering.

**Graphical Abstract:** 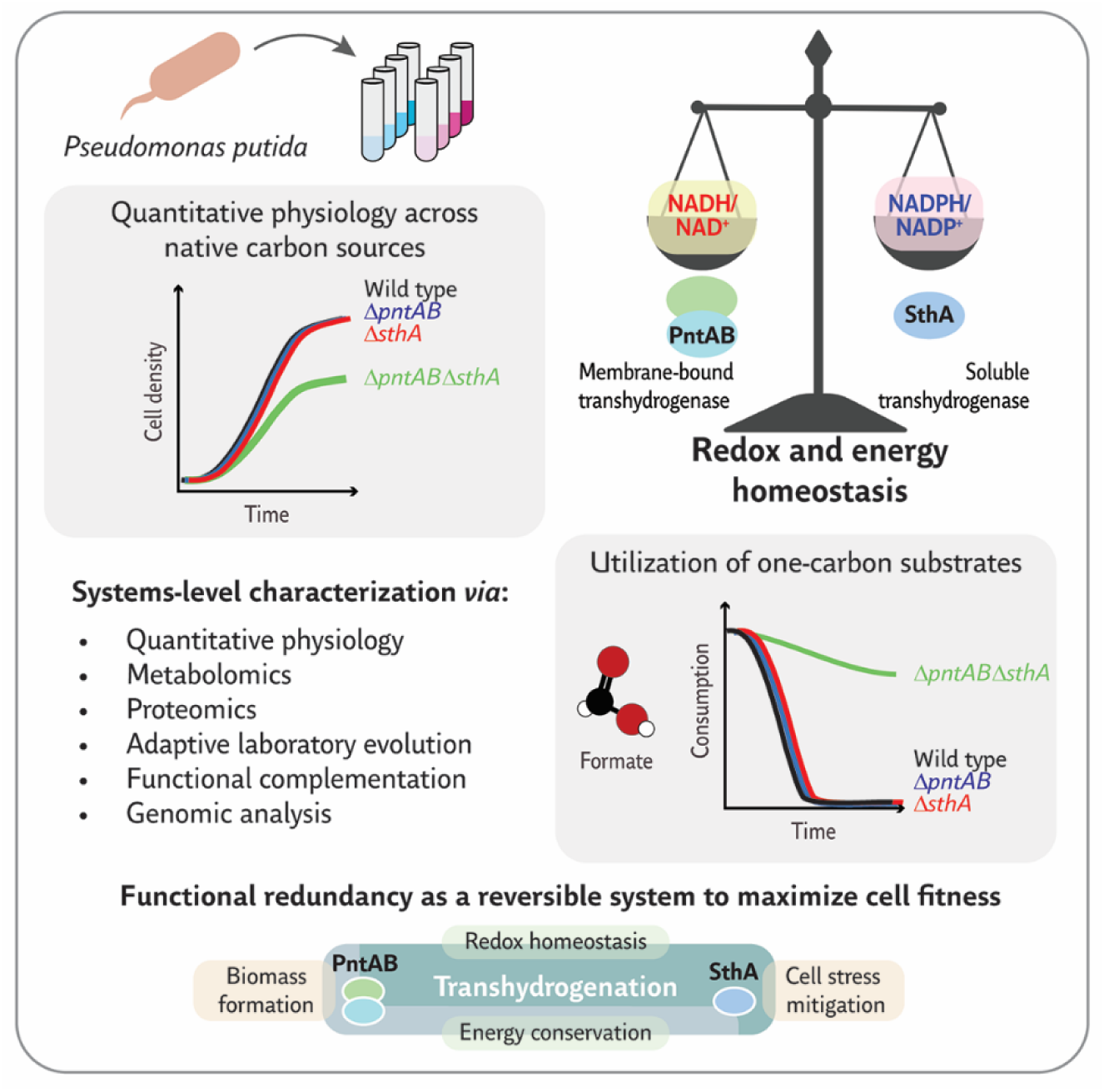

## Introduction

Redox homeostasis, essential for all living cells (Jones and Sies, 2015; Pollak et al., 2007; Shimizu and Matsuoka, 2019) drives multiple vital metabolic processes. The nicotinamide adenine dinucleotide cofactors NAD(H) and NADP(H) are conserved hub metabolites across all domains of life (Partipilo et al., 2023a; Schmidt et al., 2003) functioning as carriers of reducing equivalents. Bacteria maintain NADH/NAD^+^ and NADPH/NADP^+^ ratios within tightly defined concentration ranges through enzymatic balancing mechanisms that provide the redox potential required for supporting anabolism and catabolism (Fuhrer and Sauer, 2009; Spaans et al., 2015). Transhydrogenases are among the enzymes responsible for redox balancing, as they catalyze a hydride transfer between NAD(H) and NADP(H). Transhydrogenase enzymes belong to two types: the H^+^-translocating, membrane-bound transhydrogenase Pnt and the soluble, FAD-containing Sth.

In *Escherichia coli*, PntAB is known to primarily generate NADPH for biosynthesis, whereas SthA dissipates excess NADPH by producing NADH for oxidative phosphorylation (Sauer et al., 2004). Yet, *in vivo* and *in vitro* studies questioned the strict separation of these divergent functions. For example, *sthA* overexpression in engineered *E*. *coli* increased poly(3-hydroxybutyrate) synthesis by regenerating NADPH rather than depleting the cofactor (Sánchez et al., 2006). In *Comamonas testosteronii*, which lacks SthA, PntAB alone supports either NADH or NADPH regeneration depending on carbon source availability (Wilkes et al., 2023). *In vitro* experiments further emphasized functional flexibility. Reconstitution of PntAB in liposomes showed that hydride transfer directionality depended on the H^+^-motive force (Graf et al., 2021), whereas SthA catalyzed both NADPH- and NADH-dependent reactions (Cao et al., 2011; Partipilo et al., 2022) and was implemented for NAD(P)H regeneration in cell-free systems (Boonstra et al., 2000b; Mouri et al., 2009; Partipilo et al., 2021). This relaxed reversibility is connected with the near-identical standard reduction potentials of NAD^+^/NADH + H^+^ and NADP^+^/NADPH + H^+^ with *E*_0_’ = −0.32 V (Burton and Wilson, 1953). Together, these observations suggest that transhydrogenases play a more flexible physiological role than previously assumed.

*Pseudomonas putida* is a suitable model to explore metabolic flexibility in redox metabolism. This model soil bacterium is widely used as a synthetic biology *chassis* (Calero and Nikel, 2019; de Lorenzo et al., 2024; Volke et al., 2020a) owing, among other features, to its capacity to sustain redox-intensive reactions (Akkaya et al., 2018; Blank et al., 2008; Ebert et al., 2011; Nikel et al., 2021; Zobel et al., 2017). The elevated rates of NAD(P)H turnover (Nikel et al., 2015) are connected to the metabolic architecture of *P*. *putida* (**Fig. 1A**), which harbors both PntAB and SthA (**Fig. 1B**). The membrane-bound transhydrogenase is encoded by *pntAA* (*PP_0156*), *pntAB* (*PP_5747*), and *pntB* (*PP_0155*). Although the three genes form a single transcriptional unit, *pntAB* was only recently recognized as a functional gene (Nikel et al., 2016), which explains the non-correlative numbering of the open reading frames. Before this finding, *pntAB* was considered to be a pseudogene (Nelson et al., 2002). The soluble transhydrogenase, in contrast, is encoded by a single gene, *PP_2151*. AlphaFold-assisted structural analysis verified the membrane-bound nature of PntAB and the soluble character of SthA (**Fig. 1C**). The combined activity of PntAB and SthA is involved in supporting redox balance in *P*. *putida* during aromatic compound degradation (Nikel et al., 2016). As it happens in bacteria other than *E*. *coli* or *Bacillus subtilis*—considered as the model Gram-negative and Gram-positive organisms, respectively—the role, regulation, and flexibility of transhydrogenation in *P*. *putida* remain unexplored.

**Figure 1.**
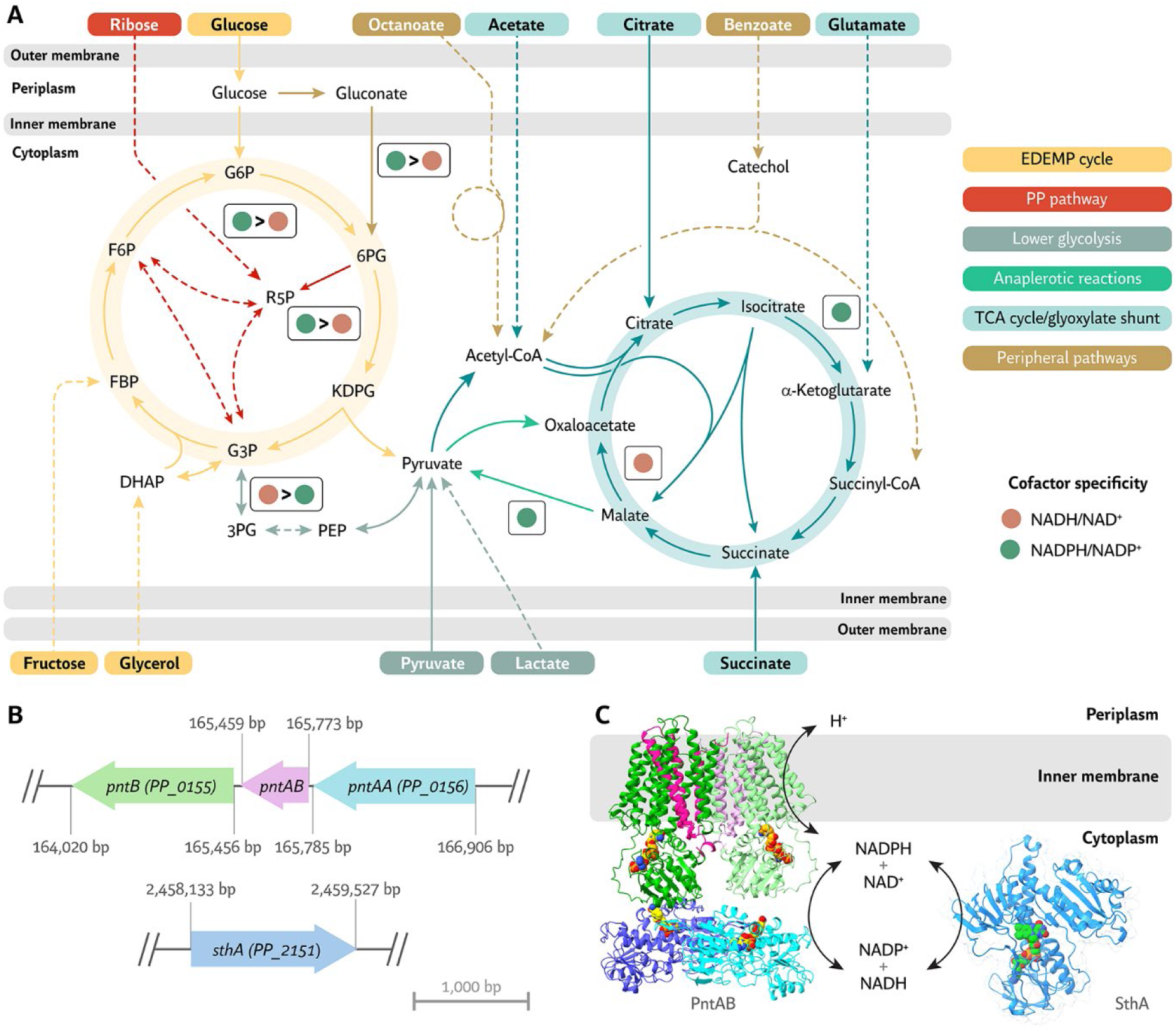
The membrane-bound and soluble transhydrogenases within the metabolic architecture of *Pseudomonas putida*. **(A)** Central carbon metabolism of *P*. *putida* and cofactor specificities for the main NAD^+^- and NADP^+^-dependent dehydrogenases. Representative carbon sources are color-coded by the main metabolic route involved in their assimilation. NADP^+^ specificity is marked with a green circle, NAD^+^ specificity with a brown circle. Dashed lines indicate multiple reactions; solid lines denote transport or single enzymatic steps. *Abbreviations:* EDEMP cycle, Entner-Doudoroff–Embden-Meyerhof-Parnas–pentose phosphate cycle; PP pathway, pentose phosphate pathway; TCA cycle, tricarboxylic acid cycle; 6PG, 6-phosphogluconate; 2-keto-3-deoxy-6-phosphogluconate; G6P, glucose-6-phosphate; F6P, fructose-6-phosphate; FBP, fructose-1,6-bisphosphate; DHAP, dihydroxyacetone phosphate; G3P, glyceraldehyde-3-phosphate; R5P, ribose-5-phosphate; 3PG, 3-phosphoglycerate; PEP, phosphoenolpyruvate; and CoA, coenzyme A. **(B)** The flavin-dependent soluble transhydrogenase is encoded by *sthA* (*PP_2151*), while the membrane-bound transhydrogenase PntAB is formed by the products of three distinct genes, *pntAA* (*PP_0156*), *pntAB* (*PP_5747*), and *pntB* (*PP_0155*), organized in a single transcriptional unit. **(C)** Structure of PntAB and SthA in *P*. *putida*. PntAB comprises a membrane-embedded domain that translocates H^+^ across the lipid bilayer, and two soluble domains that mediate hydride transfer between NADP(H) and NAD(H). In the homodimeric assembly of PntAB, PntB contains most of the membrane domain (dark and light green) and the NADP(H)-binding domain, PntAB contributes three membrane-spanning helices (magenta and pink), and PntAA corresponds to the NAD(H)-binding domain (purple and cyan). SthA is a FAD-containing soluble enzyme with an undefined oligomeric state. NAD^+^, NADP^+^, and FAD are represented as yellow, orange, and green spheres, respectively. Cofactor positioning was inferred by superimposing the AlphaFold structures of PntAA, PntAB, and PntB (Uniprot codes: A0A140FVX4, A0A140FVX3, and Q88RH4) with the structure of the transhydrogenase from *Ovis aries* (PDB code: 6QTI), or *via* AlphaFill for the predicted structure of SthA (Uniprot code: Q88KY8) using a homologous structure (PDB code: 2EQ7).

Redox balancing during metabolism of one-carbon (C_1_) feedstocks, e.g., formate and methanol, has only recently gained attention as a critical factor for implementing synthetic C_1_ assimilation pathways. In *P*. *putida*, glucose co-feeding with formate increased expression of the *pntAB* operon (Turlin et al., 2023). Transhydrogenases were also implicated in evolutionary adaptation to formate assimilation. Formatotrophic growth of an engineered *E*. *coli* rapidly improved after the strain acquired a mutation in the *pntAB* promoter that enhanced transcription by an order of magnitude (Kim et al., 2020). Similarly, placing a strong constitutive promoter to boost *pntAB* transcription supported efficient formate assimilation in engineered *P*. *putida* (Turlin et al., 2025; Turlin et al., 2022). The connection between C_1_ metabolism and transhydrogenase activity, however, remains unresolved.

Here, we systematically addressed the roles of PntAB and SthA in *P*. *putida* by testing Δ*pntAB*, Δ*sthA*, and Δ*pntAB* Δ*sthA* mutants across culture conditions. In most cases, deletion of single transhydrogenase variants did not substantially impact cell physiology, a first indication of reversible activity *in vivo*. Biochemical assays and metabolomics supported the notion of flexible directionality, with redox imbalance evidenced only in the double mutant. SthA was strictly required for growth on acetate, a dependence linked to glyoxylate metabolism that emerged through adaptive laboratory evolution (ALE). Physiological analysis also established a role of transhydrogenases in formate tolerance, leading to the identification of a native, O_2_-tolerant and NAD(P)H-independent formate dehydrogenase (FDH) as a redox compensating mechanism. We also explored the cofactor specificity of downstream detoxification routes. Furthermore, comparative genomic analyses indicated associations of SthA with lipid homeostasis and oxidative stress, while PntAB was largely linked to membrane processes, signaling, and stress responses. Taken together, these results demonstrate that the rigid directionality once attributed to transhydrogenases does not hold across bacterial species and instead emphasize their broad physiological flexibility, shaped by ecological niche and evolutionary contingency.

## Results

### A single transhydrogenase variant suffices to support cellular fitness across culture conditions

To examine the role of PntAB and SthA in *P*. *putida* KT2440, we used transhydrogenase-deleted strains (Δ*pntAB*, Δ*sthA*, and Δ*pntAB* Δ*sthA*) previously developed in our laboratory to study the redox burden associated with aromatic compound degradation (Nikel et al., 2016). In a first set of experiments, the physiology of the wild-type (WT) strain and the transhydrogenase mutants was compared in rich medium, lysogeny broth (LB), and in minimal salt medium (MSM) with different carbon sources, using cell growth and physiological parameters as proxy (**Fig. 2A**). All strains showed identical growth in LB, indicating that both transhydrogenases are dispensable for growth under nutrient-rich conditions. The growth parameters of the Δ*pntAB* and Δ*sthA* mutants also matched closely those of the WT strain when incubated in MSM with glucose as the sole carbon source (**Table S1**). In contrast, the double Δ*pntAB* Δ*sthA* mutant exhibited ca. 36% slower growth than the WT (**Fig. 2A**).

**Figure 2.**
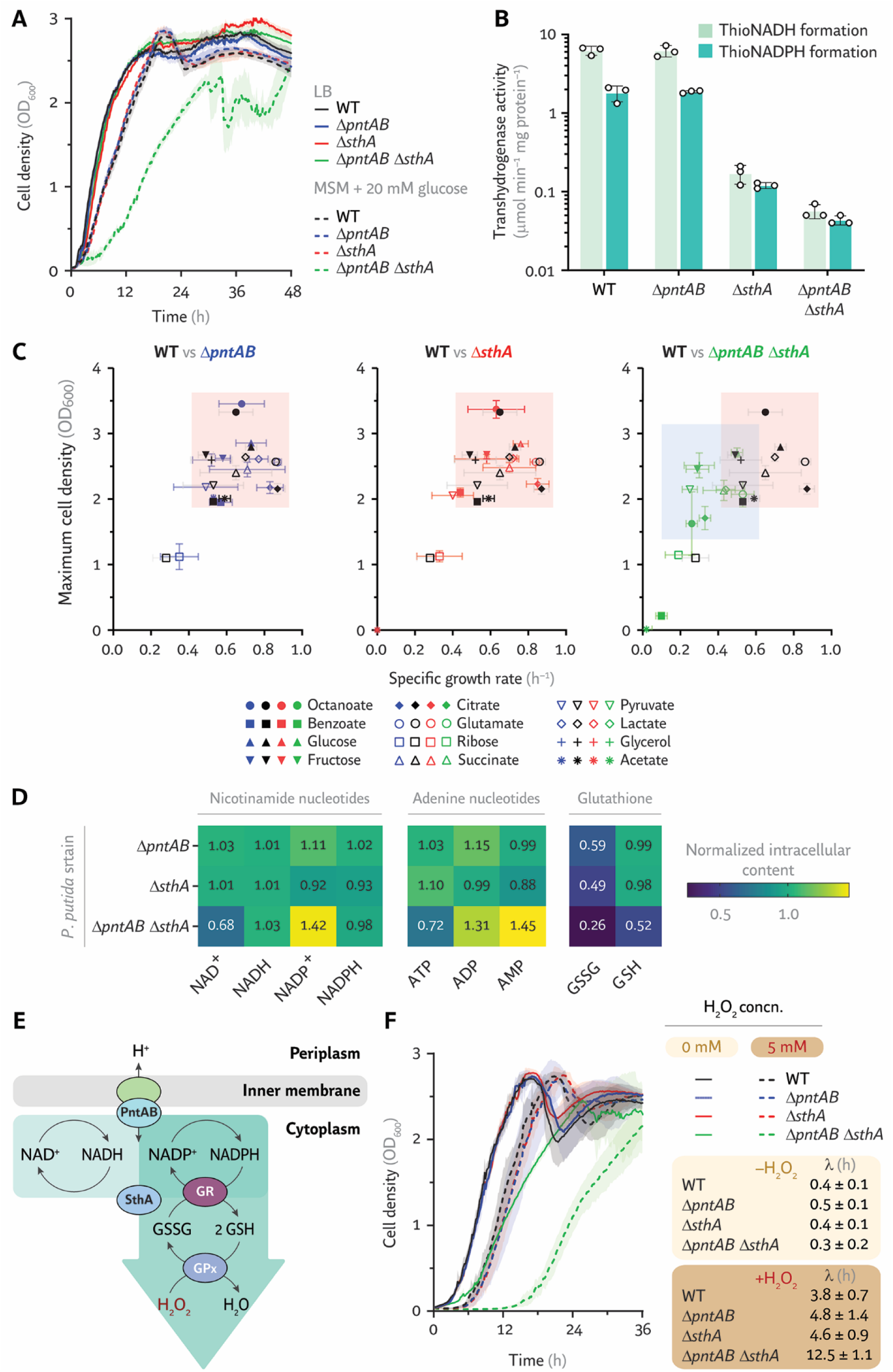
Transhydrogenases ensure optimal cellular fitness of *P*. *putida* under diverse culture conditions. **(A)** Growth of wild-type *P*. *putida* (WT) and its transhydrogenase mutant derivatives in rich and minimal media. Cell density, estimated as the optical density at 600 nm (OD_600_), was monitored in a microplate reader with either LB or minimal salt medium (MSM) containing 20 mM glucose as the sole carbon source. Experiments were performed in biological replicates (*n* = 4); data represent average values ± standard deviation. **(B)** ThioNADH- and thioNADPH-forming activities of transhydrogenases in *P*. *putida*. Reactions were carried out in cell lysates of the WT and mutant strains at 37°C (*n* = 3). Data represent average values ± standard deviation. **(C)** Impact of single and double transhydrogenase deletions on cell fitness across various carbon sources. The specific growth rate and maximum biomass density were used as proxies for fitness after a 72-h cultivation in MSM containing the substrates indicated, adjusted to provide equivalent carbon content (15 mM octanoate, 17 mM benzoate, 20 mM glucose, 20 mM fructose, 20 mM citrate, 24 mM glutamate, 24 mM ribose, 30 mM succinate, 40 mM pyruvate, 40 mM lactate, 40 mM glycerol, and 60 mM acetate). Shaded areas delineate the fitness range for the WT strain (light red) and the double mutant (light blue), excluding ribose. Experiments were performed in biological replicates (*n* = 4); data represent average values ± standard deviation. **(D)** Quantification of nicotinamide, adenosine, and glutathione metabolites in transhydrogenase mutants. Metabolite levels are shown as the intracellular concentration (μM) in the mid-exponential phase relative to WT. In the case of GSSG and GSH, the relative abundance is indicated as peak intensity at the same sampling point. Experiments were performed in biological replicates (*n* = 3). Average values are shown; errors are provided in **Fig. S2. (E)** Mechanism of transhydrogenase-dependent NADPH formation and NADPH-supported stress protection. GR, glutathione reductase; GPx, glutathione peroxidase. **(F)** Oxidative stress response of WT and mutants to H_2_O_2_ in MSM with 20 mM glucose. The extension of the lag phase (λ) is shown in the inset. Concn., concentration. Experiments were performed in biological replicates (*n* = 5); data represent average values ± standard deviation.

The unaffected growth of the single mutants provided a first suggestion on partial reversibility of transhydrogenase reactions *in vivo*, although the activity of other metabolic enzymes could also contribute to transhydrogenation. To test this possibility, directional electron transfer was measured in cell lysates using thioNAD^+^ and thioNADP^+^ as substrates (**Fig. 2B**). Thionicotinamide adenine dinucleotide and its phosphorylated form are the thione-modified derivatives of NAD^+^ and NADP^+^, respectively, and can be adopted as substrate analogues to study hydride transfer reactions (Singh et al., 2003). In the WT strain background, transhydrogenation favored NADH over NADPH formation at a 3.5-fold faster rate (6.2 ± 0.7 and 1.8 ± 0.3 μmol min^−1^ mg protein^−1^ for thioNADH and thioNADPH, respectively). Transhydrogenation rates in lysates of *P*. *putida* Δ*pntAB* were comparable to the WT, confirming that PntAB was inactive in this strain. This result also exposed that SthA could catalyze a reversible transhydrogenation reaction in the absence of PntAB. This outcome could be derived from thermodynamic considerations, although the actual transhydrogenation activity *in vivo* would be further constrained by kinetic enzyme parameters and the actual substrate concentrations. In bacteria, the NADP^+^/NADPH ratio is kept at ca. 0.8-1.5, while the NAD^+^/NADH is maintained at ca. 10-20 (Andersen and von Meyenburg, 1977; Nikel et al., 2008; Nikel et al., 2009; Schroer et al., 2009). Given that the reaction NAD^+^ + NADPH ↔ NADP^+^ + NADH has a K*_eq_*∼ 1, the reaction *in vivo* is probably kept out of equilibrium (Zhou et al., 2011), favoring the formation of NADP^+^ + NADH by the soluble transhydrogenase.

Lysates where PntAB was the only transhydrogenase (i.e., the Δ*sthA* strain) exhibited very low transhydrogenation rates, since the H^+^ gradient coupling is lost during cell disruption. Lysates of the double mutant lacked activity in both directions, confirming that no other soluble enzymes can mediate hydride transfer between NAD(P)H and NAD(P)^+^.

To further clarify the physiological role of transhydrogenases, the WT strain and mutants were grown on MSM with different carbon sources (**Table S1**). The substrates selected varied in carbon number (from C_2_ to C_8_), degree of reduction (γ), and entry point into central metabolism (**Fig. 1A**). To compare all experimental strains across conditions, cellular fitness was explored in plots of the maximum cell density (estimated as the optical density at 600 nm, OD_600_) and the specific growth rate (**Fig. 2C**; individual growth curves are shown in **Fig. S1**). Except for *P*. *putida* Δ*sthA* grown on acetate or benzoate, both single mutants maintained cellular fitness similar to the WT (**Fig. 2C**, light red shaded area). In contrast, the double mutant displayed reduced fitness on ten out of the 12 carbon sources tested (**Fig. 2C**, green data points within the blue shaded area), showing that either transhydrogenase is sufficient for supporting growth across most environmental conditions. The two clear exceptions with severely affected cellular fitness were acetate and ribose. Eliminating SthA completely abolished acetate-dependent growth. Growth on ribose was unaffected even in the double mutant, which probably reflects the naturally slow ribose-dependent growth of *P*. *putida* (Volke et al., 2021), where slower alternative mechanisms of redox balancing based on other native dehydrogenases might suffice.

To directly assess the effect of transhydrogenase deletions on cofactor balance, the intracellular content of nicotinamide cofactors (i.e., NAD^+^, NADP^+^, NADH, and NADPH), adenine nucleotides (i.e., AMP, ADP, and ATP), and glutathione species (glutathione disulfide, GSSG, and reduced glutathione, GSH) was quantified in the strains (**Fig. 2D** and **Fig. S2**). Single mutants showed no major changes in redox cofactor ratios or energy state, with all values in the same range as for the WT (**Table S2**). In contrast, *P*. *putida* Δ*pntAB* Δ*sthA* exhibited a 32% lower NAD^+^ and ATP levels compared to WT, while NADP^+^, ADP, and AMP increased by 31% to 45%. Both individual deletion strains (Δ*pntAB* and Δ*sthA*) had a decreased GSSG content (ca. 50-60% of WT), whereas the double mutant dropped to < 30%. The GSH content remained largely unchanged in the single mutants, but the double mutant displayed a 50% decrease, suggesting that neither enzyme alone provides acts as the main NADPH source but that both act synergistically to maintain the reduced glutathione pool.

The inability of the Δ*pntAB* Δ*sthA* strain to maintain stable cofactor ratios, combined with severe GSH depletion—not observed in either single mutant—led us to test whether both transhydrogenases could supply reducing power for quell oxidative stress. The main defense mechanisms against oxidative stress, glutathione reductase and glutathione peroxidase, rely on NADPH to regenerate the glutathione pool (GSSG → 2GSH) as a key intracellular redox buffer (**Fig. 2E**). To determine whether one or both transhydrogenases regenerate NADPH for this function, we compared strain growth in the presence of 5 mM H_2_O_2_ (**Fig. 2F**). Under oxidative stress conditions, cellular fitness followed the same general trends observed without the stressor. Neither SthA nor PntAB alone seemed to be preferentially required for NADPH formation for glutathione reduction, as suggested by the comparable lag phases (λ) of strains Δ*pntAB*, Δ*sthA*, and WT. In contrast, the double Δ*pntAB* Δ*sthA* mutant had a 3-fold longer λ compared to the WT strain and single mutants when exposed to H_2_O_2_, suggesting that both transhydrogenases are required to efficiently maintain the GSH pool *via* NADPH regeneration.

### The soluble SthA transhydrogenase is required for acetate assimilation *via* a regulatory mechanism

Deletion of *sthA* abolished growth of *P*. *putida* on acetate (**Fig. 2C**), raising the question of the role of the soluble transhydrogenase in assimilation of this C_2_ substrate. Proteomic analysis of the WT strain (**Fig. 3A**) showed that SthA abundance on acetate-grown cells was ca. 4-fold higher than in the glucose condition, underscoring a key role of the soluble transhydrogenase in acetate metabolism. The soluble subunits of PntAB (PntAA and PntB) were also more abundant on acetate, but their levels reached only about half of SthA, indicative of a comparatively minor contribution. Hence, to determine whether deleting *pntAB* caused broader metabolic or energetic changes in acetate cultures, the intracellular cofactor ratios were measured under these conditions (**Fig. 3B** and **Fig. S3**). The WT and Δ*pntAB* strains displayed nearly identical profiles for all parameters tested. Interestingly, cells grown on acetate showed higher NADH/NAD^+^ and NADPH/NADP^+^ ratios and a lower ATP/ADP ratio compared to glucose-grown *P*. *putida*. These findings indicate an enhanced availability of reducing power in the form of nicotinamide cofactors, independent of the activity of the membrane-bound transhydrogenase.

**Figure 3.**
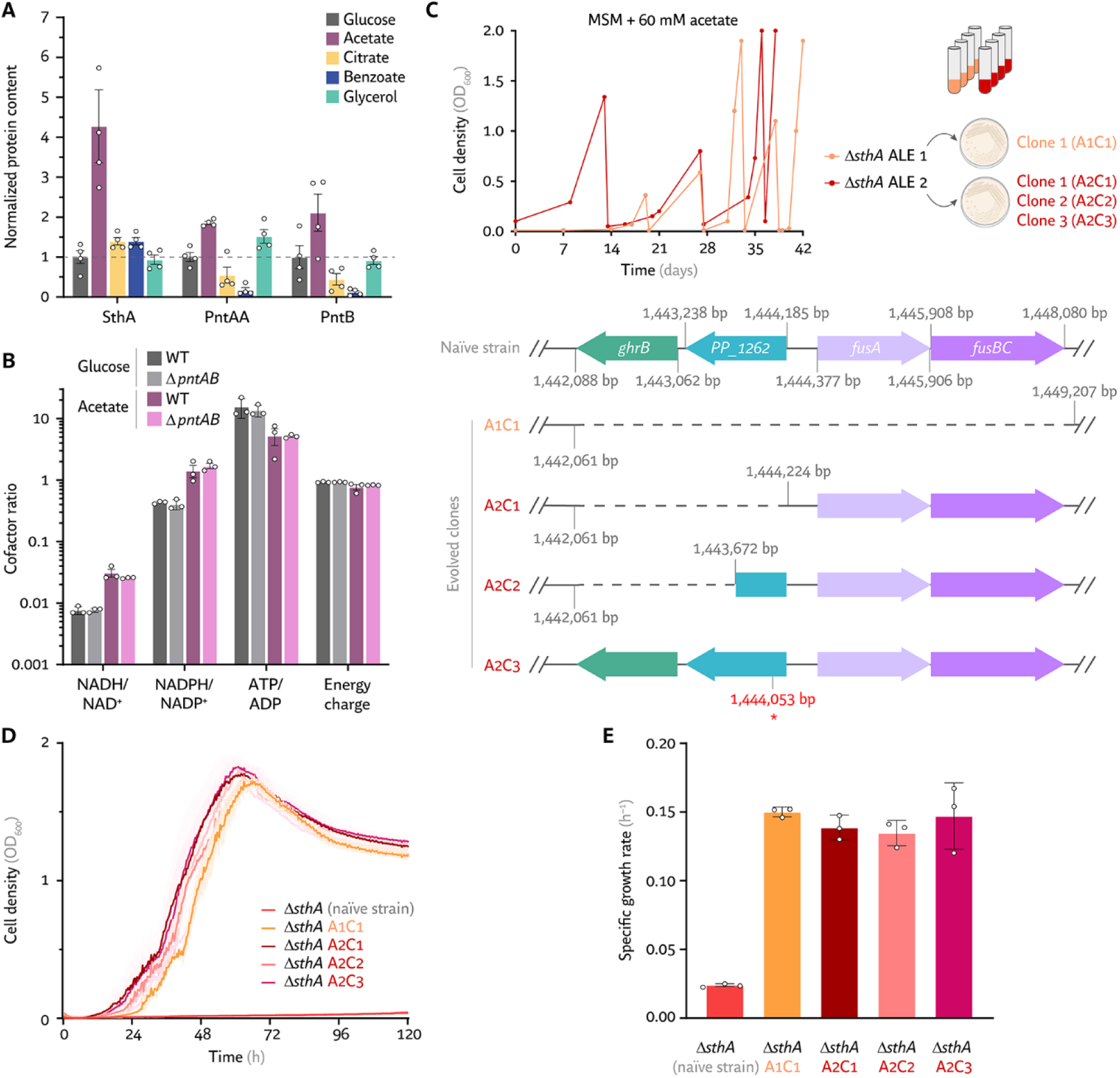
Role of the soluble transhydrogenase SthA on acetate metabolism. **(A)** Relative abundance of intracellular transhydrogenases across different carbon sources. Protein levels were normalized to the amount detected during growth in MSM containing glucose. *P*. *putida* was grown on 20 mM glucose, 20 mM acetate, 20 mM citrate, 10 mM benzoate, or 20 mM glycerol. Proteins were extracted during mid-exponential phase. Experiments were performed in biological replicates (*n* = 4); data represent average values ± standard error mean. **(B)** Cofactor ratios and energy charge in the WT and Δ*pntAB* strains during growth on glucose and acetate. Experiments were performed in biological replicates (*n* = 3); data represent average values ± standard deviation. **(C)** Adaptive laboratory evolution (ALE) restores growth of *P*. *putida* Δ*sthA* on acetate. Passage history from two independent ALE experiments is shown in the top panel. Four clones were isolated from these lines; all carried alterations within the same 6-kb region, with specific mutations for each clone shown in the bottom panel. The red asterisk denotes a point mutation introducing a premature *STOP* codon in *PP_1262* (Q45*). **(D)** Growth curves and **(E)** specific growth rates of evolved Δ*sthA* clones on acetate. Experiments were performed in biological replicates (*n* = 3); data represent average values ± standard deviation.

To expose a potential connection between *sthA* and acetate assimilation, an ALE strategy (Fernández-Cabezón et al., 2019) was employed to restore growth on acetate and identify the genetic modifications responsible for the phenotype (**Fig. 3C**). Two independent ALE experiments were performed in MSM with 60 mM acetate as the sole carbon and energy source. In both cases, cell density increased slowly over 2-3 weeks. Reinoculation of the cultures into fresh acetate-containing medium led to progressive growth improvements that culminated in biomass formation similar to the WT levels within one week. Whole-genome sequencing of the evolved strains revealed consistent alterations in a 6-kb genomic locus comprising four genes (**Fig. 3C**), encoding GhrB (PP_1261), a LysR-type transcription factor (PP_1262), and two fusaric acid resistance proteins (FusA and FusB, PP_1263 and PP_1264, respectively). The only gene disrupted in all evolved strains was the transcriptional regulator *PP_1262*, interrupted either by complete or partial deletion, as in evolved clones A1C1, A2C1, and A2C2, or by a premature *STOP* codon in clone A2C3. Growth curves and specific growth rates were similar across all evolved strain lineages (**Fig. 3D** and **3E**).

To explore a direct regulatory link between PP_1262 and GhrB, a NADP^+^-dependent 2-ketoaldonate reductase/hydroxypyruvate/glyoxylate reductase encoded by *PP_1261*, a Δ*sthA* Δ*PP_1261* double mutant was constructed and tested for growth on acetate (**Fig. S4**). Similar to the parental Δ*sthA* strain, the double mutant was unable to assimilate acetate. These findings show that the essential role of SthA in acetate metabolism in *P*. *putida* involves a regulatory mechanism dependent on the transcriptional regulator PP_1262, rather than an imbalance in reducing power. In *E*. *coli*, by contrast, the essentiality of *sthA* has been attributed to excess NADPH generation through the tricarboxylic acid (TCA) cycle (Sauer et al., 2004). While the precise function of PP_1262 remains to be defined, these results suggest a regulatory layer shaped by transhydrogenases that extends beyond redox balance.

### Synergistic role of transhydrogenase and formate dehydrogenases in formate detoxification

The results of acetate cultures prompted the question of whether tolerance to non-native substrates could also be influenced by transhydrogenase activity. Given that *P*. *putida* displays high tolerance to C_1_ feedstocks (Roca et al., 2008; Turlin et al., 2022)—a trait that supports microbial-based valorization of CO_2_-derived substrates (Favoino et al., 2025), e.g., formate (Turlin et al., 2025)—the physiological link between transhydrogenation and formate oxidation was systematically examined in cultures with mixed substrates. Formate is not a native carbon source for *P*. *putida* (**Fig. 1A**), but it can be processed through peripheral pathways that extract electrons from this C_1_ molecule. The physiological performance of the WT and transhydrogenase mutant strains was first compared in microtiter plate cultures using glucose as the main carbon source, with and without 240 mM formate, a concentration that does not halt growth while inducing stress (**Fig. 4A**). Formate supplementation reduced the specific growth rate of the WT and single-deletion mutants by ca. 35%, while the double mutant showed a stronger reduction of ca. 60%. This trend was also reflected in the normalized cell density. The Δ*pntAB* Δ*sthA* strain exhibited a ca. 70% decrease in maximum biomass when formate was supplied. In contrast, all transhydrogenase strains reached comparable maximal OD_600_ values in medium containing glucose alone. Under these conditions, no substantial improvement in biomass yield was observed, unlike the modest positive effect reported at lower formate concentrations (Zobel et al., 2017).

**Figure 4.**
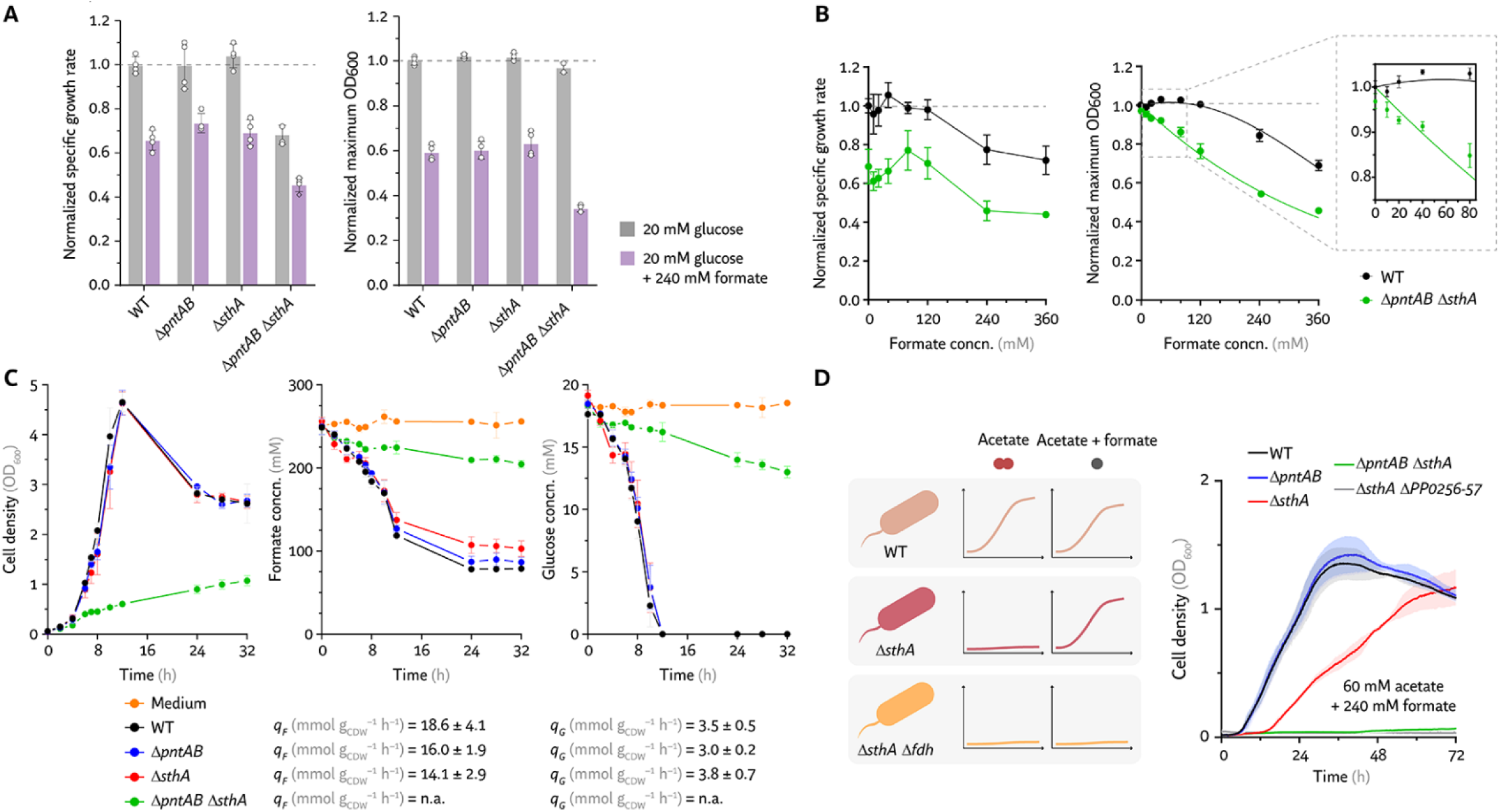
Dissecting the link between transhydrogenation and formate tolerance. **(A)** Physiological parameters of the WT strain and transhydrogenase mutants grown on glucose with or without 240 mM formate in microtiter plate cultures. Experiments were performed in biological replicates (*n* = 4); data represent average values ± standard deviation. Values were normalized to WT grown on glucose alone. **(B)** Growth response of WT and the Δ*pntAB* Δ*sthA* mutant to increasing concentrations of formate in glucose cultures. Specific growth rate (left panel) and maximum OD_600_ (right panel) were calculated and normalized to WT values for all conditions. Experiments were performed in biological replicates (*n* = 4); data represent average values ± standard deviation. **(C)** Quantitative physiology of the WT strain and the transhydrogenase mutants in shaken-flask cultures using glucose and formate. The specific rates of formate and glucose consumption (*q*_F_ and *q*_G_) were calculated from biological replicates (*n* = 3); data represent average values ± standard deviation. n.a., not applicable. **(D)** Formate oxidation restores acetate-dependent growth of the Δ*sthA* mutant. Growth of the transhydrogenase mutant, with or without the Δ*PP_0256-PP_0257* deletion, was evaluated in acetate cultures supplemented with formate in microtiter plates. Experiments were performed in biological replicates (*n* = 4); data represent average values ± standard deviation.

Formate titration in cultures of the WT and Δ*pntAB* Δ*sthA* strains provided quantitative insight into how formate affects cellular fitness (**Fig. 4B**). Growth rates decreased similarly in both strains at formate levels > 120 mM, although a slight increase (< 10%) was observed between 40 and 80 mM (**Fig. 4B**, inset). Yet, *P*. *putida* Δ*pntAB* Δ*sthA* displayed a higher sensitivity in biomass formation than the WT strain, with normalized cell densities dropping by ca. 15% at 20 mM formate and by ca. 30% at 120 mM formate. The WT strain maintained unchanged biomass levels until concentrations exceeded 120 mM.

To test whether *P*. *putida* Δ*pntAB* Δ*sthA* lost the ability to oxidize formate while metabolizing hexoses, formate and glucose consumption were measured simultaneously in shaken-flask cultures in MSM supplemented with 20 mM glucose and 240 mM formate (**Fig. 4C**). The single mutants showed virtually identical growth, formate detoxification, and glucose consumption patterns as the WT strain, supporting the largely interchangeable roles of membrane and soluble transhydrogenases in maintaining redox balance between NAD(H) and NADP(H) pools for supporting overall metabolic functions. In contrast, the double mutant displayed severely impaired formate oxidation and glucose utilization, preventing an accurate estimation of specific consumption rates (*q*) due to the absence of an authentic exponential growth phase.

While only ca. 5 mM glucose was consumed over 32 h, yielding an OD_600_ ∼1, the formate concentration declined by ca. 30 mM during the same period (**Fig. 4C**). Under equivalent culture conditions without formate, glucose consumption was comparable between the WT strain and its Δ*pntAB* Δ*sthA* derivative (**Fig. S5A** and **S5B**), substantiating that the growth and uptake defects arose exclusively in the presence of the C_1_ feedstock. Hence, deleting both transhydrogenases prevents redistribution of reducing equivalents from formate oxidation, causing redox imbalance. The inability to oxidize formate probably led to its intracellular accumulation, which disrupts carbon metabolism (Mira and Teixeira, 2013). Thus, both transhydrogenases act synergistically to channel reducing equivalents released from formate oxidation to downstream nicotinamide cofactors, sustaining redox homeostasis and growth in the presence of formate.

To expand this quantitative physiology analysis of formate-induced stress, growth was tested across multiple carbon sources. A consistent defect was observed only in *P*. *putida* Δ*pntAB* Δ*sthA*, although its severity depended on the substrate (**Fig. S6**). The strongest impairment occurred when formate was co-fed with benzoate, completely abolishing growth. This outcome likely reflects the combined effects of formate-induced oxidative stress and the high redox demand required for aromatic compound degradation (Nikel et al., 2016). Surprisingly, *P*. *putida* Δ*sthA* was able to grow on acetate when cultures were supplemented with formate, suggesting a formate-dependent mechanism that supports slow, but consistent growth on acetate (**Fig. 4D**). Under these conditions, the final OD_600_ of the Δ*sthA* strain reached comparable values to those of the WT and Δ*pntAB* mutant, although the specific growth rate was reduced by ca. 40% with λ ∼ 14 h. To investigate the basis of this compensatory mechanism, absent in glucose cultures, the enzymes directly involved in formate processing in *P*. *putida* were examined. A FDH system formed by PP_0256 and PP_0257 was recently identified in our laboratory as central for formate tolerance (Turlin et al., 2023), and we targeted this FDH to explore synergies between formate oxidation and redox balance afforded by transhydrogenases. When a Δ*PP_0256*-*PP_0257* deletion was introduced into the Δ*sthA* background, growth on acetate with 240 mM formate was completely abolished—exposing an essential role for this FDH in formate detoxification in the absence of SthA.

### Plugging-in NAD^+^- and NADP^+^-dependent FDHs reveals cofactor specificity of formate detoxification pathways

The role of the FDH encoded by *PP_0256-PP_0257* in relieving oxidative stress appeared evident when the cells were exposed to high formate concentrations, but it remained unclear whether this enzymatic route ultimately transferred the reducing equivalents from formate oxidation to NAD^+^ or NADP^+^. PP_0257 corresponds to FdhD, a sulfurtransferase required for the activation of molybdenum-dependent FDHs (Thomé et al., 2012), whereas *PP_0256* encodes a molybdopterin-binding oxidoreductase sharing 59% identity and 71% similarity with Fdh4A from *Methylobacterium extorquens* (Chistoserdova et al., 2007). Fdh4A was reported as essential for *in vivo* formate oxidation to CO_2_, but no protein-level characterization clarified its catalytic mechanism. To address this question, bioinformatic analyses were used to assess the potential requirement of nicotinamides in PP_0256-mediated catalysis. Sequence-based searches for a Rossmann fold (Geertz-Hansen et al., 2014; Kamiński et al., 2022), a structural motif typical of nicotinamide binding, failed to reveal a site specific for NAD(P)^+^ recognition. This outcome prompted a structure-guided approach based on the AlphaFold-predicted model of the PP_0256 protein (PDB entry: AF-Q88R78-F1). Substrate transplantation *via* AlphaFill (Hekkelman et al., 2023) also ruled out direct NAD(P)^+^ binding, instead showing structural similarity to the experimentally determined FDH H from *E*. *coli* (**Fig. 5A**), a component of the anaerobic formate-hydrogen lyase (FHL) (Kammel et al., 2024) complex that embeds two molybdopterin guanine dinucleotide and one Fe_4_S_4_ cluster for formate oxidation to CO_2_ (PDB entries: 1AA6 and 1FDI, respectively) (Boyington et al., 1997). These findings suggested that the reducing equivalents generated by PP_0256 during formate detoxification are transferred through other unknown protein subunits likely embedding intermediates, possibly flavins such as FMN (as in the case of the formate dehydrogenase from *Rhodobacter capsulatus*) (Radon et al., 2020), ultimately reaching the soluble NAD(P)^+^ pool. The reoxidation of the reduced nicotinamide cofactor likely depends on a transhydrogenation-based redox balancing mechanism, consistent with the data shown in **Fig. 4A-C**.

**Figure 5.**
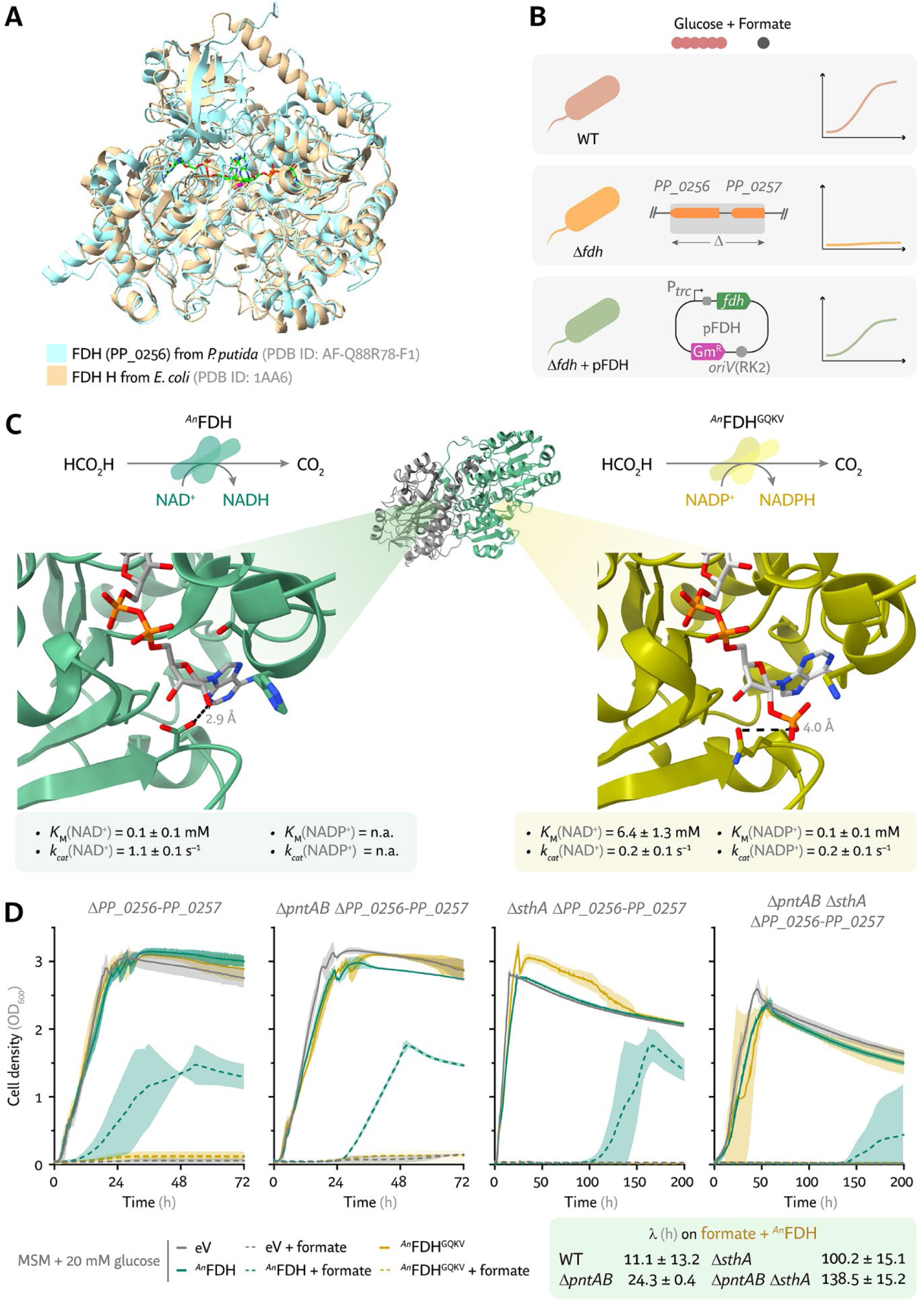
Plugging-in NAD⁺- and NADP⁺-dependent formate dehydrogenases (FDHs) to test the cofactor specificity of formate detoxification pathways. **(A)** Superimposed structures of PP_0256 from *P*. *putida* and FdhF (FDH H) from *E*. *coli*. The structure of PP_0256 was predicted using AlphaFold. **(B)** Schematic of the complementation strategy to restore formate tolerance in the formate dehydrogenase-deficient (Δ*PP_0256-PP_0257*) strain. Gm^R^, gentamicin-resistance determinant; pFDH, expression plasmid carrying different *fdh* genes. **(C)** Biochemical features of the non-metal FDH enzyme from *Ancylobacter novellus* (*^An^*FDH) and its rationally engineered variant with altered cofactor specificity (*^An^*FDH^GQKV^). n.a., not applicable. **(D)** Formate oxidation mediated by PP_0256-PP_0257 is functionally complemented by a non-metal NAD⁺-dependent FDH. Mutants of *P. putida* lacking PP_0256-PP_0257 alone or in combination with PntAB and SthA were transformed with constitutively-expressed soluble NAD⁺-dependent (*^An^*FDH) or NADP⁺-dependent (*^An^*FDH^GQKV^) formate dehydrogenase genes. The corresponding strains transformed with the empty pS621c vector, indicated as eV, were used as a negative control. Experiments were performed in biological replicates (*n* = 3); data represent average values ± standard deviation.

In an attempt to establish a link between formate oxidation and redox balancing of nicotinamide cofactors through transhydrogenation, the *PP_0256-PP_0257* genes were deleted in all transhydrogenase-mutant *P*. *putida* strains. Bacterial genes encoding non-metal FDHs with strict specificity for one nicotinamide cofactor were then expressed constitutively using the P*_trc_* promoter in vector pS621c (Turlin et al., 2023), giving rise to different pFDH plasmids (**Fig. 5B**). The NAD^+^-dependent FDH from *Ancylobacter novellus* (*^An^*FDH) and its quadruple mutant (containing the A198G, D221Q, H379K, and S380V mutations) (Partipilo et al., 2023b) with shifted cofactor preference (*^An^*FDH^GQKV^) were chosen for these experiments, due to their well-characterized biochemical and structural properties (**Fig. 5C**). In cultures containing 20 mM glucose and 240 mM formate, only the strains expressing *^An^*FDH from plasmid pFDH1 tolerated formate-induced stress. Strains harboring the empty vector pS621c or plasmid pFDH2, encoding the NADP^+^-dependent *^An^*FDH^GQKV^ variant, failed to grow within 72 h (**Fig. 5D**). This pattern held across all experimental strains, but the Δ*pntAB* mutant showed a lag phase comparable to the WT strain when expressing the NAD^+^-dependent FDH, while the Δ*sthA* strain exhibited an extended lag phase (λ) of ca. 100 h. The delay was even more pronounced in the double Δ*pntAB* Δ*sthA* mutant (λ = 138.5 ± 15.2 h). The prolonged lag observed in *P*. *putida* Δ*sthA* when formate oxidation was driven by a heterologous NAD^+^-FDH highlighted the role of the soluble transhydrogenase as the main sink for excess NADH produced by this reaction. These results also indicated that SthA catalyzes NADH oxidation efficiently and operates in a readily reversible manner. The limited capacity of PntAB to perform this role alone most likely reflects its dependence on the H^+^-motive force, which is altered when formate is fed to cultures where the native FDH system is deleted. Notably, when formate oxidation proceeded through the native PP_0256 enzyme, both Δ*pntAB* and Δ*sthA* mutants displayed similar growth patterns (**Fig. 4C**), supporting the existence of downstream intermediates mediating the transfer of reducing equivalents rather than a direct connection to the nicotinamide pool.

### Occurrence and functional significance of dual-transhydrogenase adaptation across bacterial species

The physiological roles of PntAB and SthA in *P*. *putida*, a model soil bacterium, prompted the exploration of their distribution across other organisms. To assess the broader significance of enzymatic transhydrogenation, the occurrence of PntAB and SthA sequences annotated in the UniProt database was examined in multiple species (**Fig. 6A**). The PntB subunit was used as a proxy to retrieve cases where the membrane-bound transhydrogenase was encoded in each genome. PntAB proteins were more prevalent than SthA homologs, showing ca. an order of magnitude difference in frequency. Membrane-bound transhydrogenases appeared in eukarya as mitochondrial proteins and in archaea, representing ca. 9% and 2% of the total entries, respectively, while soluble transhydrogenases were nearly absent in both domains. In bacteria, both enzymes were predominant in Pseudomonadota (i.e., 55% for PntAB and 76% for SthA) and, to a lesser extent, in Actinomycetota. In terms of relative abundance, Bacteroidota and Cyanobacteriota displayed higher PntAB occurrence, at ca. 5% and 3% respectively, whereas SthA was represented at ca. 2-4% among the Planctomycetota, Nitrospirota, and Myxococcota phyla.

**Figure 6.**
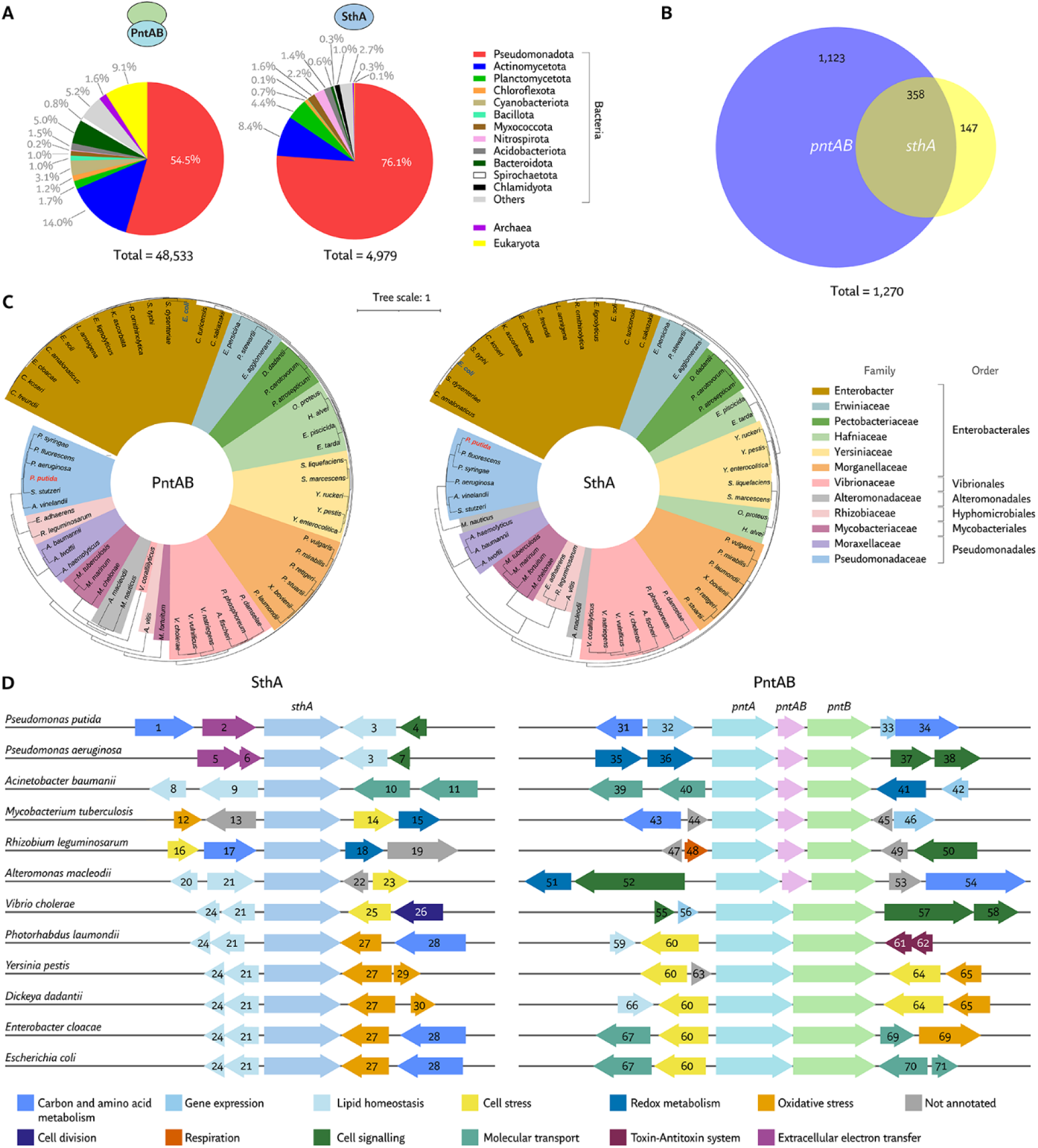
Taxonomic distribution, phylogenetic analysis, and genome context of *pntAB* and *sthA*. **(A)** Distribution of PntAB and SthA enzymes across domains of life. UniProt was queried to identify organisms with annotated homologs of both transhydrogenases. The presence of PntB was used as a proxy for the membrane-bound enzyme. Entries were grouped by taxonomic classification. **(B)** Fraction of annotated organisms harboring *pntAB* and/or *sthA* genes. The values in the Venn diagram indicate the number of organisms containing either or both transhydrogenase genes, based on the NCBI database records. **(C)** Sequence-based phylogenetic trees of bacterial transhydrogenases in representative organisms carrying both types of enzymes. The color code on the right denotes taxonomic order and family. **(D)** Genomic context diversity of *pntAB* and *sthA* across selected bacteria. The SEED database was consulted for divergent microbes to identify the functional categories of neighboring genes. Genes are numbered according to their identifiers, listed in full in **Table S5** of the Supplemental Information.

The analysis of annotated microbial genomes revealed that ca. two-thirds of bacteria harboring the *sthA* gene also contained the membrane-bound transhydrogenase genes (**Fig. 6B**). Conversely, ca. three-quarters of bacteria with *pntAB* lacked the gene for the soluble enzyme, indicating that in many species the membrane-bound transhydrogenase alone maintains redox balance between the NADH/NAD^+^ and NADPH/NADP^+^ pools. This pattern likely reflects the mechanistic coupling of PntAB-mediated transhydrogenation with the H^+^-motive force, a process essential for ATP synthesis.

Because a significant number of microorganisms encode both transhydrogenase variants, the evolutionary distance between PntAB and SthA was analyzed in representative bacteria possessing both enzymes (**Fig. 6C** and **Table S3**). Comparable phylogenetic trees derived from multiple alignments of each enzyme showed clustering according to family and taxonomic order. The mutational rates of PntAB and SthA were similar and independent of their co-occurrence. This observation mirrored the phylogenetic profile of RsmA (**Fig. S7**), a ribosomal RNA small subunit methyltransferase A essential for protein translation (Seistrup et al., 2017). The dual transhydrogenase adaptation of *P*. *putida* appeared among the most evolutionarily distant from that of *E*. *coli*, the only other bacterial species where the physiological roles of both transhydrogenases have been studied in depth.

To expand these observations, the genomic contexts of both transhydrogenases were investigated across diverse bacterial families (**Fig. 6D** and **Tables S4-S5**). A guilt-by-association-based approach (Aravind, 2000; Koonin et al., 2021) was applied to identify cellular processes potentially linked to transhydrogenation and redox homeostasis. The *sthA* gene frequently colocalized with genes involved in lipid homeostasis and oxidative stress. This feature was supported by the recurrent upstream presence of genes encoding the unsaturated fatty acid biosynthesis repressor *fabR* (gene 21, **Fig. 6D**) and the inner membrane protein *yijD* (gene 24) (Zhu et al., 2009), as well as the downstream occurrence of *oxyR* (gene 27), which encodes a transcriptional regulator of antioxidant-related gene expression (Lushchak, 2011). To a lesser extent, *sthA* was also found near genes associated with carbon and amino acid metabolism, such as argininosuccinate lyase (gene 28), and stress response functions across taxonomically diverse bacteria.

In contrast, the genomic context of the membrane-bound, H^+^-translocating transhydrogenase operon was more variable, suggesting a greater degree of evolutionary plasticity. The operon structure ranged from three distinct genes (i.e., *pntA*, *pntAB*, and *pntB*) to an overlapping two-gene configuration (i.e., *pntA* and *pntB*), suggesting that a gene fusion event occurred in an ancestral lineage followed by differential retention across taxa. Despite these structural variations, stress-related genes were often conserved near *pntAB* in several bacterial species. *Photorhabdus laumondii*, *Yersinia pestis*, *Dickeya dadantii*, *Enterobacter cloacae*, and *E*. *coli* shared the upstream presence of *ydgH* (gene 60, **Fig. 6D**) (Eletsky et al., 2014), which encodes a cell envelope stress-associated protein. In some cases, *pntAB* was flanked downstream by *uspE* (gene 64), which encodes a universal stress protein (Xu et al., 2016). Genes related to oxidative stress regulation were also detected near *pntAB* in some species, including those encoding the Fnr transcriptional regulator and a *S*-(hydroxymethyl)glutathione dehydrogenase (genes 65 and 69, respectively). Additionally, molecular transport and signaling genes often appeared within the *pntAB* genomic neighborhood in distantly related species, suggesting potential interactions between H^+^-dependent redox balancing and diverse membrane processes, e.g., nutrient uptake, efflux, and signal transduction.

## Discussion

Since the discovery of enzymatic transhydrogenation in bacterial extracts (Colowick et al., 1952), the exchange of reducing equivalents between NAD(H) and NADP(H) mediated by PntAB and SthA has intrigued microbiologists and biochemists alike. Although transhydrogenase enzymes are not universally essential, as evidenced by their absence in some bacterial lineages, they are widespread and generally occur either as a single enzyme, with the membrane-bound PntAB being more common than the soluble SthA counterpart, or as two structurally unrelated proteins coexisting in the same microorganism. This long-standing observation raises a fundamental question regarding the physiological advantage of maintaining two structurally and mechanistically distinct enzymes catalyzing the same reversible reaction (Blank et al., 2010; Voordouw et al., 1983). In *E.coli*, this distinction has been attributed to opposing directionalities *in vivo*. In this model, PntAB harnesses the H^+^-motive force to reduce NADP⁺ to NADPH using NADH as the electron donor, while SthA reoxidizes excess NADPH to generate NADH, used in the respiratory chain for energy conservation. Whether this framework applies to the phylogenetic diversity of bacteria has remained unresolved.

The investigation of dual transhydrogenase function in *P*. *putida* revealed a more flexible metabolic system. Rather than operating in settled directions, PntAB and SthA are the components of a cooperative mechanism that adjusts redox balance dynamically, a view contrasting with the traditional, rigid functional interpretation derived from model organisms. A detailed physiological characterization of Δ*pntAB* and Δ*sthA* mutants yielded several key findings that support this notion. First, the deletion of either enzyme had negligible effects on cellular fitness under diverse metabolic regimes, regardless of the carbon source entry point or its degree of reduction (**Fig. 2C** and **Table S1**). Second, intracellular concentrations of NAD(H), NADP(H), adenosine cofactors, and reduced glutathione in the mutants matched the profiles of the WT strain during growth on glucose and acetate (**Fig. 2D** and **Table S2**). Third, exposure to oxidative stress induced by H_2_O_2_ did not impair redox resilience in either mutant, which maintained NADPH-dependent antioxidant pools comparable to WT (**Fig. 2F**). Fourth, both WT and single mutants oxidized formate *via* the PP_0256-0257 FDH at nearly identical rates (**Fig. 4C**). In contrast, the double mutant Δ*pntAB* Δ*sthA* had impaired growth, energy imbalance, and heightened oxidative stress sensitivity across multiple environmental conditions. Disruption of both enzymes reduced the energy charge and altered cofactor ratios due to NAD^+^ and NADP^+^ imbalances, leading to suboptimal energy conservation and slowed metabolic fluxes. No experimental evidence supported a fixed *in vivo* directionality toward NADPH or NADH formation. Instead, both enzymes appear to be rather reversible, with the actual catalytic direction determined by intracellular kinetics and flux demands. PntAB likely favors NADPH generation under certain regimes, whereas SthA may produce NADH, but both switch roles dynamically to sustain redox homeostasis.

An exception emerged during growth on acetate, where SthA alone proved indispensable. This phenotype was linked to the LysR-type transcriptional regulator encoded by *PP_1262*. The absence of any discernable growth in Δ*sthA* on acetate resembles the observations in trashydrogenase mutant of *E. coli* (Sauer et al., 2004), though the mechanism differs. *E*. *coli* lacks a close homolog of PP_1262, and related regulators share < 37% sequence identity (**Fig. S8**). The regulatory network controlled by PP_1262 in *P*. *putida*, either directly or indirectly, remains to be fully defined.

Once the roles of PntAB and SthA were clarified on standard carbon sources, their contribution during formate co-feeding was examined. Enzymatic transhydrogenation proved essential to counter redox stress derived from formate feeding. Neither transhydrogenase acted preferentially in this process. Instead, PntAB and SthA functioned synergistically to mediate electron redistribution, enabling efficient formate detoxification and utilization, which ensured maximal biomass accumulation (**Fig. 4C**). Remarkably, formate supplementation restored growth of *P*. *putida* Δ*sthA* on acetate by overcoming PP_1262-mediated repression. Electrons from formate oxidation were likely rerouted through PntAB to support acetate metabolism (**Fig. 4D**).

The coupling between formate oxidation and transhydrogenation for redox balance was strictly dependent on PP_0256-PP_0257, the major FDH of *P*. *putida*. Structural analysis through AlphaFold modeling, artificial intelligence-guided substrate transplantation, and fold recognition revealed a strong similarity between PP_0256 and FDH H from *E*. *coli*, which oxidizes formate anaerobically without the need of soluble nicotinamide cofactors (Boyington et al., 1997). This suggests that downstream proteins, and not PP_0256 itself, transfer electrons to nicotinamide cofactors in *P*. *putida*. These yet unidentified elements appear to require the concerted action of PntAB and SthA to maintain redox equilibrium. Complementation assays with the NAD⁺-dependent *^An^*FDH substantiated that NAD⁺ serves as the ultimate electron acceptor during formate oxidation (**Fig. 5C-D**). This experimental setup uncovered an interaction between NADH generated by *^An^*FDH and SthA activity. When both *PP_0256-PP_0257* and *sthA* were deleted, the strain expressing *^An^*FDH displayed a 4-fold longer lag phase than that of the Δ*PP_0256-PP_0257* and Δ*PP_0256-PP_0257* Δ*pntAB* strains harboring the same construct. Together, these findings support the selection of *P*. *putida*, a non-C_1_-trophic organism, as a suitable host for engineering synthetic formatotrophy (Favoino et al., 2025), a promising route for circular biomanufacturing based on CO_2_-derived formate (Orsi et al., 2023; Puiggené et al., 2025a; Yishai et al., 2016). Understanding the native mechanisms driving formate oxidation will enable rational design of efficient synthetic C_1_ assimilation modules. Artificial intelligence-based and bioinformatic tools will be instrumental in this effort by identifying structural homologs inaccessible to sequence-based methods (Baltoumas et al., 2023; Cheskis et al., 2025; Partipilo and Slotboom, 2024). Along the same line of reasoning, methylotrophy has become another key direction for engineering *P*. *putida* platforms assimilating C_1_ feedstocks. Synthetic linear and cyclic pathways for methylotrophy show promise for enabling methanol assimilation in engineered *P*. *putida* strains (Puiggené et al., 2025b; Turlin et al., 2025), although the role of transhydrogenases in supporting these designs remains largely underexplored. For example, increased *pntAB* expression was essential for restoring methylotrophic growth in a *M*. *extorquens* mutant deficient in methanol utilization (Carroll and Marx, 2013). Given the diversity of metabolic and regulatory networks of natural and synthetic C_1_-trphic bacteria, transhydrogenases provide a general mechanism for channeling reducing equivalents from C_1_ feedstocks into biosynthetic routes.

The redundancy of PntAB and SthA in *P*. *putida* thus acts as a redox safety valve enabling coordinated exchange between NAD(H) and NADP(H) pools; a reversible system that maintains optimal fitness and minimizes redox stress. Genomic analysis revealed that *sthA* occurs in conserved redox-stress-adapted contexts, whereas *pntAB* is largely associated with more diverse, membrane-linked genomic loci (**Fig. 6D**). These patterns reflect distinct evolutionary trajectories of cytoplasmic, soluble *versus* membrane-bound transhydrogenases in bacterial redox homeostasis. Considering the reported directionalities for the enzymes in *E*. *coli*, the phylogenetic divergence separating *P*. *putida* from Enterobacteria (**Fig. 6C**) could explain, to an extent, distinct functional adaptations. Supporting this notion, SthA in Pseudomonadaceae differs structurally from the octameric form found in *E*. *coli*. In *Pseudomonas fluorescens* and *Azotobacter vinelandii*, SthA forms extended filaments composed of repeating subunits hundreds of nanometers long (Boonstra et al., 2000a; Boonstra et al., 1999; French et al., 1997; Shen et al., 2025). Such filaments are often subject to allosteric regulation, and in SthA from *A*. *vinelandii*, disassembly is triggered by NADP^+^ (Lynch et al., 2020; Park and Horton, 2019; van den Broek et al., 1971). These structural variations reflect divergent regulatory mechanisms *in vivo*, explaining the contrasting catalytic directionalities between *E*. *coli* and *P*. *putida*. Further *in vitro* studies could be designed to compare kinetic and regulatory properties of both transhydrogenases across species. Examining dual systems in bacteria progressively separated phylogenetically may reveal how evolutionary pressures and redox demands shaped the diverse functional landscapes of bacterial transhydrogenation. Such insights will be critical for understanding the general principles governing redox metabolism, while supporting the development of synthetic metabolism that exploits transhydrogenase versatility for biotechnology.

In summary, this study establishes that transhydrogenases coordinate redox and energy metabolism in *P*. *putida*, extending their role beyond peripheral redox valves. These enzymes are essential to sustain fitness during formate detoxification by fine-tuning balancing NAD(H) and NADP(H) pools. The coupling of formate oxidation to transhydrogenation enables redistribution of reducing equivalents toward biosynthesis, stress resistance, and energy conservation (**Fig. 7**). This network-wide integration of diverse cellular processes positions transhydrogenases as active modulators of metabolic fluxes, connecting peripheral detoxification with central carbon metabolism. Thus, the dual system of soluble and membrane-bound isoforms supports electron flow redistribution across conditions, contributing to the metabolic robustness typical of soil bacteria. As such, these findings outline a mechanistic basis for redox control in *P*. *putida* and inform engineering strategies targeting cofactor-dependent pathways—especially towards synthetic assimilation of C_1_ substrates, which relies on native or engineered redox metabolism (Orsi et al., 2024).

**Figure 7.**
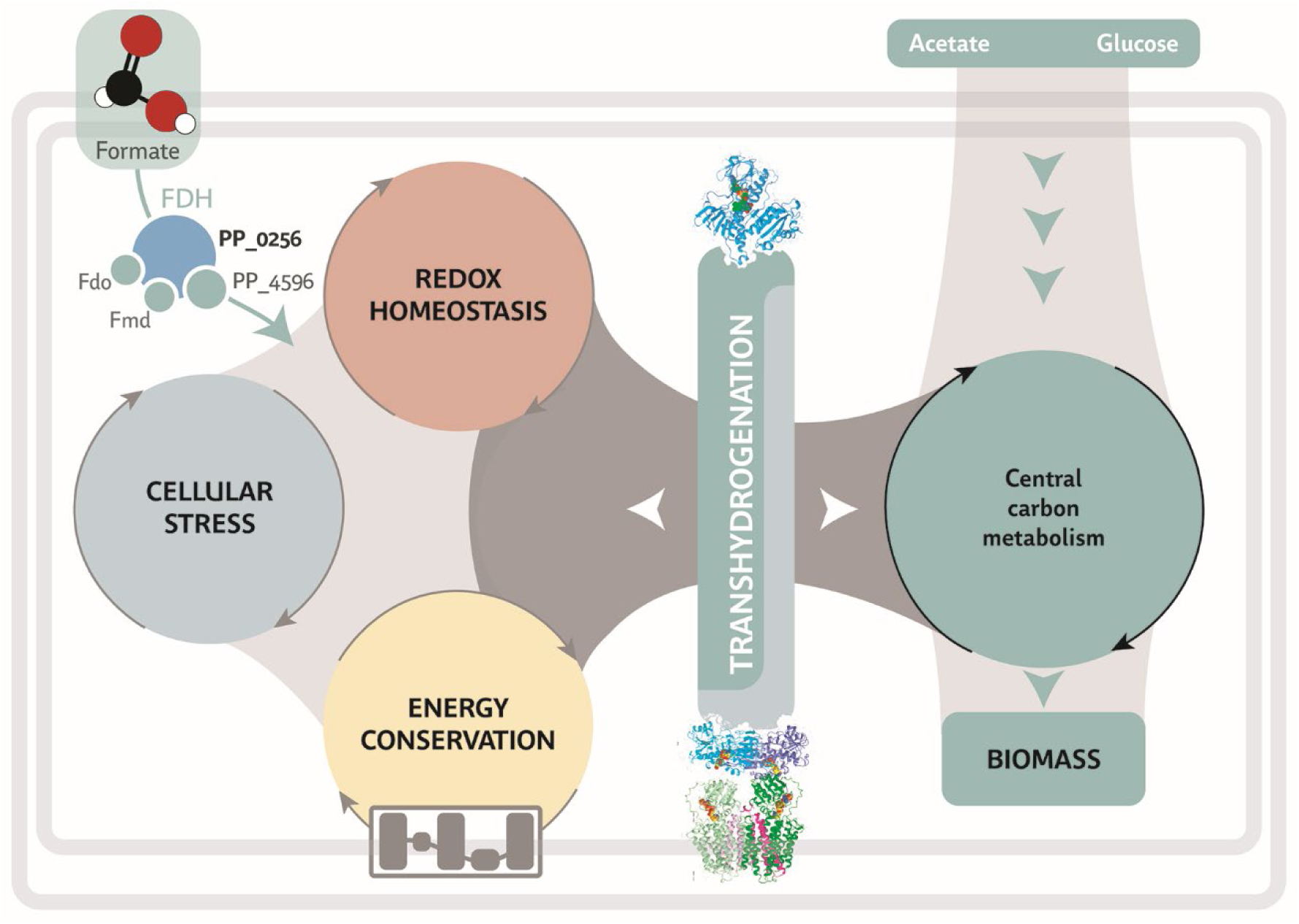
Role of transhydrogenases in integrating central carbon metabolism with formate detoxification in *Pseudomonas putida*. Oxidation of formate by the NAD^+^-dependent formate dehydrogenases PP_0256 and PP_4596 contributes to redox input into the metabolic network. The two transhydrogenases of *P*. *putida* mediate electron transfer between the NAD(H) and NADP(H) pools, linking redox homeostasis, energy conservation, and the cellular stress response. This reversible interconversion ensures that reducing equivalents generated during formate oxidation are redistributed to meet the cofactor demands of biosynthesis and oxidative stress defense. Thus, transhydrogenation serves as an interface between peripheral detoxification routes and central carbon metabolism, enabling network-wide flux balance.

## Methods

### Bacterial strains and cultivation conditions

Bacterial strains, plasmids, and oligonucleotides used in this study are listed in **Tables S6-S8**. *P. putida* KT2440 and its derivative deletion strains served as the main models for quantitative physiology experiments, while *E*. *coli* DH5α λ*pir* was used as the cloning host. The Δ*pntAB*, Δ*sthA*, and Δ*pntAB* Δ*sthA* strains previously generated (Nikel et al., 2016) were tested to confirm the presence of the corresponding deletions by colony PCR (**Fig. S9**).

All strains were cultivated in lysogeny broth (LB) or minimal salt medium (MSM), both prepared as described previously (Turlin et al., 2023) at 30°C and 250 rpm. In MSM, a specific carbon source was supplied depending on the experiment, ensuring that 120 mM of total carbon atoms was present in the final medium. The following concentrations were used: 15 mM sodium octanoate, 17 mM sodium benzoate, 20 mM glucose, fructose, or sodium citrate, 24 mM sodium glutamate or ribose, 30 mM disodium succinate, and 40 mM sodium lactate, sodium pyruvate, or glycerol. For acetate experiments, 60 mM sodium acetate was provided. The effect of formate on bacterial growth and physiology parameters was evaluated by supplementing the medium with 240 mM sodium formate. Oxidative stress was induced by adding 5 mM H_2_O_2_ to MSM containing glucose.

Growth assays were conducted in an Epoch2 microtiter plate reader (BioTek Instruments Inc., Winooski, VT, USA) using 150 μL of cell suspension at an initial OD_600_ ∼ 0.05, covered with 50 μL of sterile mineral oil, or in baffled Erlenmeyer flasks under the same initial optical density. Shaken-flask cultures were grown in 40 mL of the corresponding culture medium in 250-mL Erlenmeyer flasks at 30°C and 250 rpm. For selection of plasmids and strains during genetic manipulations, kanamycin and gentamicin were added at final concentrations of 50 μg mL^−1^ and 10 μg mL^−1^, respectively. Cultures for metabolomics were carried out in 20-mL Pioreactors (Pioreactor Inc., Waterloo, Canada) operated in batch mode at 30°C with magnetic stirring at 1,000 rpm. Online measurements of OD_600_ and specific growth rates were recorded during the entire experiment. In these experiments, all *P*. *putida* strains were grown in MSM with either 20 mM glucose or 60 mM acetate as the carbon source.

### *In vitro* enzyme activities

Cell lysates were prepared from 10-mL cultures grown at 30°C and 250 rpm in MSM supplemented with 20 mM glucose as the sole carbon source. Three independent batches of cultures were used to obtain biological triplicates for enzymatic testing. When OD_600_ reached values between 0.9 and 1.1, cells were harvested by centrifugation at 4,000×*g* for 8 min at 4°C, washed once with 50 mM Tris·HCl buffer at pH = 8.0, resuspended in 1 mL of the same buffer, and disrupted by glass bead vortexing (425-600 μm glass beads; Sigma-Aldrich Co., St. Louis, MO, USA; cat. # G87721) using six alternating cycles of 1 min vortexing and 1 min resting on ice. After lysis, cell debris was removed by centrifugation at 13,000×*g* for 3 min at 4 °C. Total protein concentration in each lysate was quantified in a microplate reader by interpolation against known standards of Pierce bovine serum albumin (Thermo Fisher Scientific, Waltham, MA, USA; cat. # 23209) using the Bradford Reagent (Sigma-Aldrich Co., cat. # B6916) according to the manufacturer’s protocol.

Transhydrogenase activity in lysate samples was measured at 400 nm using thio-analogues of NAD⁺ and NADP⁺ to distinguish the reduction of thionicotinamide (surrogate) cofactors from interference by native reduced cofactors. Enzymatic assays were conducted by loading 180 μL of lysate into the wells of a microplate reader, incubating for 5 min at 37°C, and initiating the reaction by adding 20 μL of substrate mixture. The mixtures contained either 20 mM thioNAD^+^ and 10 mM NADPH to monitor thioNADH formation (final concentrations 1 mM thioNAD^+^ and 0.5 mM NADPH in a 200-μL reaction volume) or 10 mM thioNADP^+^ and 40 mM NADH to monitor thioNADPH formation (final concentrations 0.5 mM thioNADP^+^ and 2.0 mM NADH in a 200-μL reaction volume). Reactions were carried out at 37°C for 30 min in an Epoch2 microtiter plate reader, with absorbance at 400 nm recorded every 1 min. The concentrations of generated thioNADH and thioNADPH were calculated using the Lambert-Beer law and their respective extinction coefficients (ε^thioNADH^ at 400 nm = 11.9 mM^−1^ cm^−1^ and ε^thioNADPH^ at 400 nm = 11.7 mM^−1^ cm^−1^) (Anderson et al., 1963) estimating the light path length from the absorbance values of Milli-Q water at 977 nm and 900 nm.

### Metabolomics

At the point of maximum growth rate, corresponding to the mid-exponential phase, 1-3 mL of culture was collected depending on the measured OD_600_ value. Each sample was filtered under vacuum through a 0.45-μm PVDF membrane, and the filter was immersed in 2 mL of cold extraction buffer [2:2:1 acetonitrile:methanol:H_2_O with 0.1% (w/v) formic acid]. The samples were incubated at −20°C overnight, centrifuged at 17,000×*g* for 10 min at 0°C and concentrated. The dried extracts were reconstituted in LC-MS–grade water, and a ^13^C internal standard was added to generate isotope-diluted samples for analysis.

Samples were analyzed by LC-MS/MS (Rocha et al., 2025). LC separation was carried out on a Sciex Exion LC system (AB Sciex LLC, Framingham, MA, USA) equipped with an Acquity UPLC HSS T3 column (100 mm×2.1 mm, 1.8 μm; Waters Corp., Milford, MA, USA). Two mobile phases were used: mobile phase A, water containing 10 mM tributylamine, 10 mM acetic acid, 2% (v/v) isopropanol, and 5% (v/v) methanol; and mobile phase B, 100% isopropanol. The chromatographic gradient was defined as 0, 0, 0.5; 2.5, 0, 0.5; 4.3, 2, 0.5; 4.5, 6, 0.5; 5.5, 6, 0.5; 5.8, 11, 0.5; 6.5, 11, 0.5; 7.5, 28, 0.5; 8, 53, 0.2; 10.8, 53, 0.2; 11, 0, 0.2; 13, 0, 0.5; 16 stop [time (min), eluent B (% v/v), flow rate (mL min^−1^)]. MS analysis was performed in negative ion mode. Samples were acquired using multiple reaction monitoring, and extracted ion chromatograms (XIC) were processed with the Sciex OS 3.4.5 software.

Metabolite quantification was performed by isotope-dilution mass spectrometry using the ^13^C internal standard method; the ^12^C:^13^C abundance ratio was calculated for each metabolite. Calibration curves were established with analytical standards spiked with ^13^C internal standard for ATP, ADP, AMP, NAD⁺, NADH, NADP⁺, and NADPH. The linear regression range was defined as concentrations deviating < 50% from the maximum slope. Under these conditions, all calibration curves had *R*^2^ values > 0.99. Metabolite abundances (i.e., the height ratios for GSSG and GSH) and concentrations (i.e., intracellular content in μM for ATP, ADP, AMP, NAD⁺, NADH, NADP⁺, and NADPH) were normalized to the OD_600_ at the sampling point. The adenylate energy charge ratio (AEC) was calculated as 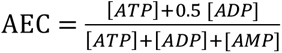.

### Sample preparation, LC-MS/MS parameters, and data acquisition for proteomics

Single colonies of *P*. *putida* KT2440 were used to inoculate overnight precultures in MSM containing different carbon sources (20 mM glucose, 20 mM potassium acetate, 20 mM glycerol, 10 mM sodium benzoate, and 20 mM sodium citrate). A 24-well Millicell Cell Culture Plate (Merck, obtained *via* Thermo Fisher Scientific Inc.) was prepared for inoculation of the overnight cultures in fresh medium, with each well containing 1 mL of the respective culture medium. Bacterial growth was monitored using a BioTek Synergy H1 microtiter plate reader at 30°C with shaking at 200 rpm, recording OD_600_ every 15 min. When the cultures reached the mid-exponential phase, cells were harvested by centrifugation at 10,000×*g* for 10 min at 4°C. The supernatant was discarded and the resulting pellets were frozen for proteomic sample preparation as described previously (Gurdo et al., 2023). Cell pellets were lysed in a buffer containing 6 M guanidinium hydrochloride, 5 mM *tris*(2-carboxyethyl)phosphine, 10 mM 2-chloroacetamide, and 100 mM Tris·HCl at pH = 8.5. The lysates were disrupted mechanically and subjected to heat treatment at 95°C. After centrifugation, the resulting cell-free lysates were diluted in 50 mM ammonium bicarbonate, and the protein concentration in each sample was determined using the bicinchoninic acid assay.

Proteins (20 μg per sample) were digested with a trypsin and LysC enzyme mix (Promega Corp., Madison, WI, USA) for 8 h. Digestion was stopped by adding trifluoroacetic acid, and peptides were desalted using C_18_ resin [Empore (3M), CDS Analytical, St. Paul, MN, USA] before high-performance liquid chromatography-mass spectrometry (HPLC-MS) analysis. HPLC-MS was performed using an Orbitrap Exploris 480 mass spectrometer (Thermo Fisher Scientific Inc.) coupled to an EASY-nLC 1200 HPLC system (Thermo Fisher Scientific Inc.). Peptide samples (500 ng each) were first trapped on a 2-cm C_18_ column (Thermo Fisher Scientific Inc., cat. # 164946) and then separated using a 70-min gradient from 8% to 48% (v/v) acetonitrile in 0.1% (v/v) formic acid on a 15-cm C_18_ reverse-phase analytical column (EASY-Spray^TM^ ES904) at a flow rate of 250 nL min^−1^. The mass spectrometer was operated in data-independent acquisition (DIA) mode, using the HRMS1 method described elsewhere (Xuan et al., 2020). The analysis was performed with a FAIMS Pro Interface (Thermo Fisher Scientific Inc.) employing a compensation voltage of –45 V, with additional modifications as indicated. Full MS1 spectra were acquired at a resolution of 120,000 within a scan range of 400-1,000 *m/z*, with an automatic gain control (AGC) target of 300 or automatically set maximum injection time. MS2 spectra were acquired at a resolution of 60,000, with an AGC target of 1,000 or auto-set maximum injection time and a collision energy of 32. Each cycle consisted of three DIA experiments covering a range of 200 *m/z* with a 6 *m/z* window size and 1 *m/z* overlap, with a full MS scan recorded between experiments. Raw DIA data were processed using DIA-NN with a library-free approach (Demichev et al., 2020). The analysis used smart profiling and heuristic protein inference with a false discovery rate threshold of 1%. *In silico* digestion included variable modifications such as *N*-terminal acetylation and methionine oxidation. Cleavage specificity was set to trypsin/P ("K*,R*"). Further settings included spectral library generation, FASTA search, predictor function, and activation of the match-between-runs (MBR) feature. Precursor and fragment exclusion parameters were defined as follows: fragment *m/z* range of 200-1,800, peptide length between 7 and 30 amino acids, precursor *m/z* range of 300-1,800, precursor charge between +1 and +4, and a maximum of one missed cleavage.

Before protein inference using the LFAQ algorithm (Chang et al., 2019), precursor intensities were summed based on identical sequences to derive peptide intensities filtered with an FDR threshold of 1%, similar to the import function of the aLFQ R-package (Rosenberger et al., 2014). A custom Python script interfaced with the LFAQ algorithm *via* the command line. Input data were processed separately for each sample, and the resulting outputs were merged. For downstream analyses, only protein intensity values were used. To meet the LFAQ algorithm requirements for protein concentration values, an auxiliary input file containing randomly selected protein identifiers with assigned random concentrations was generated; this supplementary input did not influence the calculated protein intensities.

Protein sequence identification was carried out by querying the reference proteome of *P*. *putida* (proteome ID: UP000000556) (Belda et al., 2016). Absolute quantification was achieved using the standard-free total protein approach (TPA) (Wiśniewski et al., 2014), implemented in a custom Python script for the quantification workflow. The calculated protein intensities from the inference algorithm were used as input for the analysis. Errors represent standard error means, and the obtained values on different carbon sources were normalized to the amount of each protein (SthA, PntAA or PntB) measured in cultures with glucose.

### Adaptive laboratory evolution

Two independent ALE experiments were conducted in MSM containing 60 mM acetate using the Δ*sthA* strain. ALE # 1 started with an initial OD_600_ ∼ 0.05 and was terminated after 48 days; ALE # 2 began with an OD_600_ ∼ 0.1 and lasted for 38 days. In both experiments, Pyrex glass test tubes containing 4 mL of medium were used throughout the procedure. Culture passages were performed manually when OD_600_ reached the maximum value, as determined with an Implen OD_600_ device (IMPLEN GmbH, München, Germany). At the end of both experiments, the final cultures were plated on LB agar, and one colony from ALE # 1 and three colonies from ALE # 2 were randomly picked and selected for whole genome sequencing. These clones were identified as AxCy, where *x* indicates the ALE experiment, and *y* identifies independently selected clones. Genomic DNA was extracted using the DNeasy Blood and Tissue Kit (Qiagen GmbH, Hilden, NRW, Germany) and quantified with a Qubit 2.0 fluorometer (Thermo Fisher Scientific Co.). The material was then sent for nanopore sequencing performed by Plasmidsaurus Inc. (Eugene, OR, USA). The raw reads were aligned to the genome of *P*. *putida* KT2440 (GenBank NC_002947) (Belda et al., 2016) using Geneious software (version 2025.1.2; Dotmatics, Boston, MA, USA) with the Minimap 2.24 mapper (Li, 2018), trimming 30 bp from both the 5′- and 3′-ends. Variations, including single nucleotide polymorphisms and InDels, were detected in Geneious, and variant calling was performed using a frequency threshold of 0.25 across all samples.

### Gene deletion procedures

Targeted deletions of the *PP_0256-PP_0257* locus were generated through homologous recombination using a plasmid derived from the suicide plasmid pGNW (Turlin et al., 2023; Wirth et al., 2020). Deletion plasmids were introduced into *P*. *putida* by conjugation. For this step, an overnight culture of the recipient *P*. *putida* strain was mixed with *E*. *coli* DH5α λ*pir* carrying plasmid pGNW·Δ*PP_0256-PP_0257* and the helper *E*. *coli* HB101 strain bearing the helper plasmid pRK600 on an LB agar plate, followed by overnight incubation. The resulting cell mixture was streaked on selective LB agar plates with 50 μg mL^−1^ streptomycin and 25 μg mL^−1^ 5-chloro-(2,4-dichlorophenoxy)phenol (Irgasan^TM^) (Federici et al., 2025). Conjugation plates were incubated at 30°C overnight, and positive transconjugants were selected. These colonies were then transformed by electroporation with the helper plasmid pQURE6·H (Volke et al., 2020b), using a Gene Pulser XCell electroporator (Bio-Rad Laboratories Inc., Hercules, CA, USA) set at 2.5 kV, 25 μF capacitance, and 200 Ω resistance in a 2-mm gap cuvette. After electroporation, the cells were recovered in 1 mL of LB supplemented with 1 mM 3-methylbenzoate (3-*m*Bz) and incubated at 30°C for 2 h. The culture was plated on LB agar containing the appropriate antibiotics and 1 mM 3-*m*Bz to induce plasmid replication and I-*Sce*I expression. Positive clones were verified by colony PCR using the One*Taq*™ master mix (New England Biolabs, Ipswich, MA, USA). The pQURE6·H plasmid was subsequently cured from the deletion strains by culturing cells in LB without 3-*m*Bz and selective antibiotics through several passages. The same procedure was applied to delete gene *PP_1261* using plasmid pSNW·Δ*PP_1261*, constructed *via* USER cloning as previously described (Calero et al., 2022). Primer_011 through _014 are the oligonucleotides used for vector assembly, and primer_015 and primer_016 were used to confirm the successful completion of the deletion procedure (**Table S8**).

### Analytical procedures for quantitative physiology experiments

Glucose and formate concentrations were measured in culture supernatants and analyzed using a Dionex UltiMate 3000 HPLC (Thermo Fisher Scientific Inc.) equipped with an Aminex HPX-87H column (7.8 × 300 mm, 9 μm; Bio-Rad Laboratories Inc.) and a refractive index detector Shodex RI-101 (Showa Denko America Inc., Huntington Beach, CA, USA). The column was maintained at 50°C, with a running time of 30 min and using 5 mM H_2_SO_4_ as mobile phase. Raw HPLC data were processed using the Chromeleon^TM^ 7.1.3 software package (Thermo Fisher Scientific Inc.), and metabolite concentrations were calculated from peak areas using calibration curves, each run with five standard concentrations. Specific consumption rates for glucose (*q*_G_) and formate (*q*_F_) were calculated using the following formula:

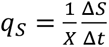

where *q_S_* represents the biomass specific substrate consumption rate, *X* is the average biomass concentration between two sampling time points, Δ*S* is the difference in substrate concentration between two sampling time points, and Δ*t* refers to the time between two sampling points.

### Construction of expression plasmids

Plasmid pS621c (Turlin et al., 2023), used as a control vector in this study, was selected as the template for constitutive expression of the non-metal formate dehydrogenase (*Snov_3272*) from *Ancylobacter novellus* DSM 506 (Partipilo et al., 2023b), in order to heterologously produce either the WT version (*^An^*FDH, NAD^+^-FDH) or a quadruple mutant (*^An^*FDH^GQKV^) that displays switched cofactor specificity (NADP^+^-FDH). The genes encoding NAD^+^-FDH and NADP^+^-FDH were PCR-amplified from expression vectors pBXC3H_*synfdh* and pBXC3H_*synfdh*^A198GD221QH379KS380V^, respectively, using the same primer pair designed for the USER cloning procedure (primer_001 and primer_002). The pS621c backbone was amplified with primer_003 and primer_004. PCR amplification was carried out using the Phusion *U* Hot Start DNA mix (Thermo Fisher Scientific Inc.), followed by treatment with 1 μL of USER^TM^ enzyme (New England BioLabs) and 1 μL of DpnI in a total reaction volume of 12 μL. The reaction contained both the amplified backbone (50 ng) and the gene of interest at a molar ratio of 1:10. USER mixtures were incubated for 30 min at 37°C, followed by a gradual temperature decrease from 28°C to 20°C over 3 min (2°C per step) and a final incubation at 10°C for 30 min. Five microliters of the USER mixtures were then transformed into chemically competent *E*. *coli* DH5α λ*pir* cells and plated on LB agar supplemented with 10 ng μL^−1^ gentamicin. Sequencing confirmed the correct assembly of the plasmids pFDH1 (encoding *^An^*FDH) and pFDH2 (encoding *^An^*FDH^GQKV^). The verified constructs were subsequently introduced into both WT and transhydrogenase-mutant strains of *P*. *putida* by electroporation. This procedure was performed in a 2-mm gap cuvette using a Gene Pulser XCell electroporator (Bio-Rad Laboratories) with settings of 2.5 kV, 25 μF capacitance, and 200 Ω resistance. Successful transformants were selected on LB agar plates containing gentamicin and were used for subsequent experimental procedures.

### Taxonomic distribution of genes and proteins

The UniProt database (release 2025_01) (Consortium, 2024) was queried using the entries «*pntB transhydrogenase*» and «*sthA transhydrogenase*» to retrieve all annotated sequences corresponding to the membrane-bound and soluble transhydrogenases, respectively. The retrieved sequences were grouped by taxonomical domains and phyla using the “taxonomy” section of the database. The Venn diagram displaying the fractions of organisms harboring the genes encoding PntAB and SthA was generated with the web tool Venny v. 2.1, available at https://bioinfogp.cnb.csic.es/tools/venny/. The corresponding datasets were extracted from the public NCBI gene database (https://www.ncbi.nlm.nih.gov/gene/) using the same search entries. Lists of organisms carrying *pntAB* and *sthA* were analyzed as previously described to identify bacteria that possess both genes in their genomes (Volke et al., 2021). Data were also curated manually for accuracy. Representative microorganisms exhibiting dual transhydrogenase adaptation were selected (**Table S3**), and the amino acid sequences of PntB and SthA homologs were retrieved from UniProt for multisequence alignment and phylogenetic analysis *via* ClustalOmega (Sievers et al., 2011). The resulting phylogenetic trees were visualized and annotated using the Interactive Tree Of Life (iTOL) v. 7 online tool (Letunic and Bork, 2021). The SEED database (Overbeek et al., 2014) (available at https://pubseed.theseed.org) was used to analyze the genomic context of *sthA* and *pntAB* in various bacterial species. Among the bacterial genomes encoding both transhydrogenases, 12 representative genomic contexts were selected for graphical representation (**Table S4**). After identifying *sthA* and *pntAB* within the genomes of interest using the SEED Viewer, the corresponding coding sequences were examined. Functional annotations of adjacent genes were retrieved to evaluate potential operon structures and metabolic associations (**Table S5**).

## Acknowledgements

The Nikel Lab acknowledges financial support from The Novo Nordisk Foundation through grants NNF20CC0035580, *LiFe* (NNF18OC0034818), *TARGET* (NNF21OC0067996), FM·*Pseudomonas* (NNF24OC0091501), and *NovoF* (NNF23OC0083631), and the European Union’s Horizon2020 Research and Innovation Programme under grant agreements Nos. 814418 (*SinFonia*) and 101082049 (*TOLERATE*) to P.I.N., and the Novo Nordisk Foundation Copenhagen Bioscience Ph.D. program grant (NNF20CC0035596) to G.F. C.M. acknowledges support from Joachim Herz Foundation and International Graduate School of Science and Engineering, TUM, and S.D. acknowledges support from the Novo Nordisk Foundation (NNF20CC0035580).

## References

Akkaya, Ö., Pérez-Pantoja, D., Calles, B., Nikel, P. I., de Lorenzo, V., 2018. The metabolic redox regime of *Pseudomonas putida* tunes its evolvability toward novel xenobiotic substrates. mBio. 9, e01512–01518.

Andersen, K. B., von Meyenburg, K., 1977. Charges of nicotinamide adenine nucleotides and adenylate energy charge as regulatory parameters of the metabolism in *Escherichia coli*. J. Biol. Chem. 252, 4151–4156.

Anderson, B. M., Anderson, C. D., Lee, J. K., Stein, A. M., 1963. The thionicotinamide analogs of DPN and TPN. ii. Enzyme studies. Biochemistry. 2, 1017–1022.

Aravind, L., 2000. Guilt by association: contextual information in genome analysis. Genome Res. 10, 1074–1077.

Baltoumas, F. A., Karatzas, E., Páez-Espino, D., Venetsianou, N. K., Aplakidou, E., Oulas, A., Finn, R. D., Ovchinnikov, S., Pafilis, E., Kyrpides, N. C., Pavlopoulos, G. A., 2023. Exploring microbial functional biodiversity at the protein family level—from metagenomic sequence reads to annotated protein clusters. Front. Bioinform. 3, 1157956.

Belda, E., van Heck, R. G. A., López-Sánchez, M. J., Cruveiller, S., Barbe, V., Fraser, C., Klenk, H. P., Petersen, J., Morgat, A., Nikel, P. I., Vallenet, D., Rouy, Z., Sekowska, A., Martins dos Santos, V. A. P., de Lorenzo, V., Danchin, A., Médigue, C., 2016. The revisited genome of *Pseudomonas putida* KT2440 enlightens its value as a robust metabolic *chassis*. Environ. Microbiol. 18, 3403–3424.

Blank, L. M., Ebert, B. E., Buehler, K., Bühler, B., 2010. Redox biocatalysis and metabolism: molecular mechanisms and metabolic network analysis. Antiox. Red. Signal. 13, 349– 394.

Blank, L. M., Ionidis, G., Ebert, B. E., Bühler, B., Schmid, A., 2008. Metabolic response of *Pseudomonas putida* during redox biocatalysis in the presence of a second octanol phase. FEBS J. 275, 5173–5190.

Boonstra, B., Björklund, L., French, C. E., Wainwright, I., Bruce, N. C., 2000a. Cloning of the sth gene from *Azotobacter vinelandii* and construction of chimeric soluble pyridine nucleotide transhydrogenases. FEMS Microbiol. Lett. 191, 87–93.

Boonstra, B., French, C. E., Wainwright, I., Bruce, N. C., 1999. The *udhA* gene of *Escherichia coli* encodes a soluble pyridine nucleotide transhydrogenase. J. Bacteriol. 181, 1030– 1034.

Boonstra, B., Rathbone, D. A., French, C. E., Walker, E. H., Bruce, N. C., 2000b. Cofactor regeneration by a soluble pyridine nucleotide transhydrogenase for biological production of hydromorphone. Appl. Environ. Microbiol. 66, 5161–5166.

Boyington, J. C., Gladyshev, V. N., Khangulov, S. V., Stadtman, T. C., Sun, P. D., 1997. Crystal structure of formate dehydrogenase H: catalysis involving Mo, molybdopterin, selenocysteine, and an Fe4S4 cluster. Science. 275, 1305–1308.

Burton, K., Wilson, T. H., 1953. The free-energy changes for the reduction of diphosphopyridine nucleotide and the dehydrogenation of L-malate and L-glycerol 1-phosphate. Biochem. J. 54, 86–94.

Calero, P., Gurdo, N., Nikel, P. I., 2022. Role of the CrcB transporter of *Pseudomonas putida* in the multi-level stress response elicited by mineral fluoride. Environ. Microbiol. 24, 5082–5104.

Calero, P., Nikel, P. I., 2019. Chasing bacterial *chassis* for metabolic engineering: a perspective review from classical to non-traditional microorganisms. Microb. Biotechnol. 12, 98–124.

Cao, Z., Song, P., Xu, Q., Su, R., Zhu, G., 2011. Overexpression and biochemical characterization of soluble pyridine nucleotide transhydrogenase from *Escherichia coli*. FEMS Microbiol. Lett. 320, 9–14.

Carroll, S. M., Marx, C. J., 2013. Evolution after introduction of a novel metabolic pathway consistently leads to restoration of wild-type physiology. PLoS Genet. 9, e1003427.

Chang, C., Gao, Z., Ying, W., Fu, Y., Zhao, Y., Wu, S., Li, M., Wang, G., Qian, X., Zhu, Y., He, F., 2019. LFAQ: toward unbiased label-free absolute protein quantification by predicting peptide quantitative factors. Anal. Chem. 91, 1335–1343.

Cheskis, S., Akerman, A., Levy, A., 2025. Deciphering bacterial protein functions with innovative computational methods. Trends Microbiol. 33, 434–446.

Chistoserdova, L., Crowther, G. J., Vorholt, J. A., Skovran, E., Portais, J. C., Lidstrom, M. E., 2007. Identification of a fourth formate dehydrogenase in *Methylobacterium extorquens* AM1 and confirmation of the essential role of formate oxidation in methylotrophy. J. Bacteriol. 189, 9076–9081.

Colowick, S. P., Kaplan, N. O., Neufeld, E. F., Ciotti, M. M., 1952. Pyridine nucleotide transhydrogenase. I. Indirect evidence for the reaction and purification of the enzyme. J. Biol. Chem. 195, 95–105.

Consortium, T. U., 2024. UniProt: the Universal Protein knowledgebase in 2025. Nucleic Acids Res. 53, D609–D617.

de Lorenzo, V., Pérez-Pantoja, D., Nikel, P. I., 2024. *Pseudomonas putida* KT2440: the long journey of a soil-dweller to become a synthetic biology *chassis*. J. Bacteriol. 206, e00136–00124.

Demichev, V., Messner, C. B., Vernardis, S. I., Lilley, K. S., Ralser, M., 2020. DIA-NN: Neural networks and interference correction enable deep proteome coverage in high throughput. Nat. Methods. 17, 41–44.

Ebert, B. E., Kurth, F., Grund, M., Blank, L. M., Schmid, A., 2011. Response of *Pseudomonas putida* KT2440 to increased NADH and ATP demand. Appl. Environ. Microbiol. 77, 6597–6605.

Eletsky, A., Michalska, K., Houliston, S., Zhang, Q., Daily, M. D., Xu, X., Cui, H., Yee, A., Lemak, A., Wu, B., Garcia, M., Burnet, M. C., Meyer, K. M., Aryal, U. K., Sanchez, O., Ansong, C., Xiao, R., Acton, T. B., Adkins, J. N., Montelione, G. T., Joachimiak, A., Arrowsmith, C. H., Savchenko, A., Szyperski, T., Cort, J. R., 2014. Structural and functional characterization of DUF1471 domains of *Salmonella* proteins SrfN, YdgH/SssB, and YahO. PLoS One. 9, e101787.

Favoino, G., Puiggené, Ò., Nikel, P. I., 2025. A blueprint for designing the next-generation of synthetic C_1_ microbes. Nat. Commun. 16, 8843.

Federici, F., Luppino, F., Aguilar-Vilar, C., Mazaraki, M. E., Petersen, L. B., Ahonen, L., Nikel, P. I., 2025. CIFR (*Clone-Integrate-Flip-out-Repeat*): a toolset for iterative genome and pathway engineering of Gram-negative bacteria. Metab. Eng. 88, 180–195.

Fernández-Cabezón, L., Cros, A., Nikel, P. I., 2019. Evolutionary approaches for engineering industrially-relevant phenotypes in bacterial cell factories. Biotechnol. J. 14, 1800439.

French, C. E., Boonstra, B., Bufton, K. A., Bruce, N. C., 1997. Cloning, sequence, and properties of the soluble pyridine nucleotide transhydrogenase of *Pseudomonas fluorescens*. J. Bacteriol. 179, 2761–2765.

Fuhrer, T., Sauer, U., 2009. Different biochemical mechanisms ensure network-wide balancing of reducing equivalents in microbial metabolism. J. Bacteriol. 191, 2112–2121.

Geertz-Hansen, H. M., Blom, N., Feist, A. M., Brunak, S., Petersen, T. N., 2014. Cofactory: sequence-based prediction of cofactor specificity of Rossmann folds. Proteins. 82, 1819–1828.

Graf, S. S., Hong, S., Müller, P., Gennis, R., von Ballmoos, C., 2021. Energy transfer between the nicotinamide nucleotide transhydrogenase and ATP synthase of *Escherichia coli*. Sci. Rep. 11, 21234.

Gurdo, N., Taylor Parkins, S. K., Fricano, M., Wulff, T., Nielsen, L. K., Nikel, P. I., 2023. Protocol for absolute quantification of proteins in Gram-negative bacteria based on QconCAT-based labeled peptides. STAR Protoc. 4, 102060.

Hekkelman, M. L., de Vries, I., Joosten, R. P., Perrakis, A., 2023. AlphaFill: enriching AlphaFold models with ligands and cofactors. Nat. Methods. 20, 205–213.

Jones, D. P., Sies, H., 2015. The redox code. Antioxid. Redox Signal. 23, 734–746.

Kamiński, K., Ludwiczak, J., Jasiński, M., Bukala, A., Madaj, R., Szczepaniak, K., Dunin-Horkawicz, S., 2022. Rossmann-toolbox: a deep learning-based protocol for the prediction and design of cofactor specificity in Rossmann fold proteins. Brief Bioinform. 23, bbab371.

Kammel, M., Erdmann, C., Sawers, R. G., 2024. The formate-hydrogen axis and its impact on the physiology of enterobacterial fermentation. Adv. Microb. Physiol. 84, 51–82.

Kim, S., Lindner, S. N., Aslan, S., Yishai, O., Wenk, S., Schann, K., Bar-Even, A., 2020. Growth of *E*. *coli* on formate and methanol *via* the reductive glycine pathway. Nat. Chem. Biol. 16, 538–545.

Koonin, E. V., Makarova, K. S., Wolf, Y. I., 2021. Evolution of microbial genomics: conceptual shifts over a quarter century. Trends Microbiol. 29, 582–592.

Letunic, I., Bork, P., 2021. Interactive Tree Of Life (*iTOL*) v5: an online tool for phylogenetic tree display and annotation. Nucleic Acids Res. 49, W293–W296.

Li, H., 2018. Minimap2: pairwise alignment for nucleotide sequences. Bioinformatics. 34, 3094–3100.

Lushchak, V. I., 2011. Adaptive response to oxidative stress: bacteria, fungi, plants and animals. Comp. Biochem. Physiol. C Toxicol. Pharmacol. 153, 175–190.

Lynch, E. M., Kollman, J. M., Webb, B. A., 2020. Filament formation by metabolic enzymes— a new twist on regulation. Curr. Opin. Cell Biol. 66, 28–33.

Mira, N. P., Teixeira, M. C., 2013. Microbial mechanisms of tolerance to weak acid stress. Front. Microbiol. 4, 416.

Mouri, T., Shimizu, T., Kamiya, N., Goto, M., Ichinose, H., 2009. Design of a cytochrome P450BM3 reaction system linked by two-step cofactor regeneration catalyzed by a soluble transhydrogenase and glycerol dehydrogenase. Biotechnol. Prog. 25, 1372– 1378.

Nelson, K. E., Weinel, C., Paulsen, I. T., Dodson, R. J., Hilbert, H., Martins dos Santos, V. A. P., Fouts, D. E., Gill, S. R., Pop, M., Holmes, M., Brinkac, L., Beanan, M., DeBoy, R. T., Daugherty, S., Kolonay, J., Madupu, R., Nelson, W., White, O., Peterson, J., Khouri, H., Hance, I., Chris Lee, P., Holtzapple, E., Scanlan, D., Tran, K., Moazzez, A., Utterback, T., Rizzo, M., Lee, K., Kosack, D., Moestl, D., Wedler, H., Lauber, J., Stjepandic, D., Hoheisel, J., Straetz, M., Heim, S., Kiewitz, C., Eisen, J. A., Timmis, K. N., Düsterhöft, A., Tümmler, B., Fraser, C. M., 2002. Complete genome sequence and comparative analysis of the metabolically versatile *Pseudomonas putida* KT2440. Environ. Microbiol. 4, 799–808.

Nikel, P. I., Chavarría, M., Fuhrer, T., Sauer, U., de Lorenzo, V., 2015. *Pseudomonas putida* KT2440 strain metabolizes glucose through a cycle formed by enzymes of the Entner-Doudoroff, Embden-Meyerhof-Parnas, and pentose phosphate pathways. J. Biol. Chem. 290, 25920–25932.

Nikel, P. I., Fuhrer, T., Chavarría, M., Sánchez-Pascuala, A., Sauer, U., de Lorenzo, V., 2021. Reconfiguration of metabolic fluxes in *Pseudomonas putida* as a response to sub-lethal oxidative stress. ISME J. 15, 1751–1766.

Nikel, P. I., Pérez-Pantoja, D., de Lorenzo, V., 2016. Pyridine nucleotide transhydrogenases enable redox balance of *Pseudomonas putida* during biodegradation of aromatic compounds. Environ. Microbiol. 18, 3565–3582.

Nikel, P. I., Pettinari, M. J., Ramírez, M. C., Galvagno, M. A., Méndez, B. S., 2008. *Escherichia coli arcA* mutants: metabolic profile characterization of microaerobic cultures using glycerol as a carbon source. J. Mol. Microbiol. Biotechnol. 15, 48–54.

Nikel, P. I., Zhu, J., San, K. Y., Méndez, B. S., Bennett, G. N., 2009. Metabolic flux analysis of *Escherichia coli creB* and *arcA* mutants reveals shared control of carbon catabolism under microaerobic growth conditions. J. Bacteriol. 191, 5538–5548.

Orsi, E., Hernández-Sancho, J. M., Remeijer, M. S., Kruis, A. J., Volke, D. C., Claassens, N. J., Paul, C. E., Bruggeman, F. J., Weusthuis, R. A., Nikel, P. I., 2024. Harnessing noncanonical redox cofactors to advance synthetic assimilation of one-carbon feedstocks. Curr. Opin. Biotechnol. 90, 103195.

Orsi, E., Nikel, P. I., Nielsen, L. K., Donati, S., 2023. Synergistic investigation of natural and synthetic C1-trophic microorganisms to foster a circular carbon economy. Nat. Commun. 14, 6673.

Overbeek, R., Olson, R., Pusch, G. D., Olsen, G. J., Davis, J. J., Disz, T., Edwards, R. A., Gerdes, S., Parrello, B., Shukla, M., Vonstein, V., Wattam, A. R., Xia, F., Stevens, R., 2014. The SEED and the Rapid Annotation of microbial genomes using Subsystems Technology (RAST). Nucleic Acids Res. 42, D206–D214.

Park, C. K., Horton, N. C., 2019. Structures, functions, and mechanisms of filament forming enzymes: a renaissance of enzyme filamentation. Biophys. Rev. 11, 927–994.

Partipilo, M., Claassens, N. J., Slotboom, D. J., 2023a. A hitchhiker’s guide to supplying enzymatic reducing power into synthetic cells. ACS Synth. Biol. 12, 947–962.

Partipilo, M., Ewins, E. J., Frallicciardi, J., Robinson, T., Poolman, B., Slotboom, D. J., 2021. Minimal pathway for the regeneration of redox cofactors. JACS Au. 1, 2280–2293.

Partipilo, M., Slotboom, D. J., 2024. The *S*-component fold: a link between bacterial transporters and receptors. Commun. Biol. 7, 610.

Partipilo, M., Whittaker, J. J., Pontillo, N., Coenradij, J., Herrmann, A., Guskov, A., Slotboom, D. J., 2023b. Biochemical and structural insight into the chemical resistance and cofactor specificity of the formate dehydrogenase from *Starkeya novella*. FEBS J. 290, 4238–4255.

Partipilo, M., Yang, G., Mascotti, M. L., Wijma, H. J., Slotboom, D. J., Fraaije, M. W., 2022. A conserved sequence motif in the *Escherichia coli* soluble FAD-containing pyridine nucleotide transhydrogenase is important for reaction efficiency. J. Biol. Chem. 298, 102304.

Pollak, N., Dölle, C., Ziegler, M., 2007. The power to reduce: pyridine nucleotides—Small molecules with a multitude of functions. Biochem. J. 402, 205–218.

Puiggené, Ò., Favoino, G., Federici, F., Partipilo, M., Orsi, E., Alván-Vargas, M. V. G., Hernández-Sancho, J. M., Dekker, N. K., Ørsted, E. C., Bozkurt, E. U., Grassi, S., Martí-Pagés, J., Volke, D. C., Nikel, P. I., 2025a. Seven critical challenges in synthetic one-carbon assimilation and their potential solutions. FEMS Microbiol. Rev. 49, fuaf011.

Puiggené, Ò., Muñoz-Triviño, J., Civil-Ferrer, L., Gille, L., Schulz-Mirbach, H., Bergen, D., Erb, T. J., Ebert, B. E., Nikel, P. I., 2025b. Systematic engineering of synthetic serine cycles in *Pseudomonas putida* uncovers emergent topologies for methanol assimilation. Trends Biotechnol. 43, 2539–2565.

Radon, C., Mittelstädt, G., Duffus, B. R., Bürger, J., Hartmann, T., Mielke, T., Teutloff, C., Leimkühler, S., Wendler, P., 2020. Cryo-EM structures reveal intricate Fe-S cluster arrangement and charging in Rhodobacter capsulatus formate dehydrogenase. Nat. Commun. 11, 1912.

Roca, A., Rodríguez-Herva, J. J., Duque, E., Ramos, J. L., 2008. Physiological responses of *Pseudomonas putida* to formaldehyde during detoxification. Microb. Biotechnol. 1, 158–169.

Rocha, C., Pinto, S. P., Jensen, S. I., Nielsen, L. K., Donati, S., 2025. A robust isotope ratio LC-MS/MS workflow for high-throughput metabolic profiling of bacteria. bioRxiv. 2025.2008.2008.669109.

Rosenberger, G., Ludwig, C., Röst, H. L., Aebersold, R., Malmström, L., 2014. aLFQ: an R-package for estimating absolute protein quantities from label-free LC-MS/MS proteomics data. Bioinformatics. 30, 2511–2513.

Sánchez, A. M., Andrews, J., Hussein, I., Bennett, G. N., San, K. Y., 2006. Effect of overexpression of a soluble pyridine nucleotide transhydrogenase (UdhA) on the production of poly(3-hydroxybutyrate) in *Escherichia coli*. Biotechnol. Prog. 22, 420– 425.

Sauer, U., Canonaco, F., Heri, S., Perrenoud, A., Fischer, E., 2004. The soluble and membrane-bound transhydrogenases UdhA and PntAB have divergent functions in NADPH metabolism of *Escherichia coli*. J. Biol. Chem. 279, 6613–6619.

Schmidt, S., Sunyaev, S., Bork, P., Dandekar, T., 2003. Metabolites: a helping hand for pathway evolution? Trends Biochem. Sci. 28, 336–341.

Schroer, K., Zelic, B., Oldiges, M., Lütz, S., 2009. Metabolomics for biotransformations: Intracellular redox cofactor analysis and enzyme kinetics offer insight into whole cell processes. Biotechnol. Bioeng. 104, 251–260.

Seistrup, K. H., Rose, S., Birkedal, U., Nielsen, H., Huber, H., Douthwaite, S., 2017. Bypassing rRNA methylation by RsmA/Dim1during ribosome maturation in the hyperthermophilic archaeon *Nanoarchaeum equitans*. Nucleic Acids Res. 45, 2007–2015.

Shen, Y., Maggiolo, A. O., Zhang, T., Warmack, R. A., 2025. CryoEM-enabled visual proteomics reveals de novo structures of oligomeric protein complexes. Structure. 33, 1484–1497.

Shimizu, K., Matsuoka, Y., 2019. Redox rebalance against genetic perturbations and modulation of central carbon metabolism by the oxidative stress regulation. Biotechnol. Adv. 37, 107441.

Sievers, F., Wilm, A., Dineen, D., Gibson, T. J., Karplus, K., Li, W., López, R., McWilliam, H., Remmert, M., Söding, J., Thompson, J. D., Higgins, D. G., 2011. Fast, scalable generation of high-quality protein multiple sequence alignments using Clustal Omega. Mol. Syst. Biol. 7, 539.

Singh, A., Venning, J. D., Quirk, P. G., van Boxel, G. I., Rodrigues, D. J., White, S. A., Jackson, J. B., 2003. Interactions between transhydrogenase and thio-nicotinamide analogues of NAD(H) and NADP(H) underline the importance of nucleotide conformational changes in coupling to proton translocation. J. Biol. Chem. 278, 33208–33216.

Spaans, S. K., Weusthuis, R. A., van der Oost, J., Kengen, S. W., 2015. NADPH-generating systems in bacteria and archaea. Front. Microbiol. 6, 742.

Thomé, R., Gust, A., Toci, R., Mendel, R., Bittner, F., Magalon, A., Walburger, A., 2012. A sulfurtransferase is essential for activity of formate dehydrogenases in *Escherichia coli*. J. Biol. Chem. 287, 4671–4678.

Turlin, J., Alván-Vargas, M. V. G., Puiggené, Ò., Donati, S., Wenk, S., Nikel, P. I., 2025. Synthetic C_1_ metabolism in *Pseudomonas putida* enables strict formatotrophy and methylotrophy *via* the reductive glycine pathway. mBio. 16, 01976–01925.

Turlin, J., Dronsella, B., De Maria, A., Lindner, S. N., Nikel, P. I., 2022. Integrated rational and evolutionary engineering of genome-reduced *Pseudomonas putida* strains promotes synthetic formate assimilation. Metab. Eng. 74, 191–205.

Turlin, J., Puiggené, Ò., Donati, S., Wirth, N. T., Nikel, P. I., 2023. Core and auxiliary functions of one-carbon metabolism in *Pseudomonas putida* exposed by a systems-level analysis of transcriptional and physiological responses. mSystems. 8, e00004–00023.

van den Broek, W. J., Santema, J. S., Wassink, J. H., Veeger, C., 1971. Pyridine-nucleotide transhydrogenase. 1. Isolation, purification and characterisation of the transhydrogenase from *Azotobacter vinelandii*. Eur. J. Biochem. 24, 31–45.

Volke, D. C., Calero, P., Nikel, P. I., 2020a. Pseudomonas putida. Trends Microbiol. 28, 512– 513.

Volke, D. C., Friis, L., Wirth, N. T., Turlin, J., Nikel, P. I., 2020b. Synthetic control of plasmid replication enables target- and self-curing of vectors and expedites genome engineering of *Pseudomonas putida*. Metab. Eng. Commun. 10, e00126.

Volke, D. C., Olavarría, K., Nikel, P. I., 2021. Cofactor specificity of glucose-6-phosphate dehydrogenase isozymes in *Pseudomonas putida* reveals a general principle underlying glycolytic strategies in bacteria. mSystems. 6, e00014–00021.

Voordouw, G., van der Vies, S. M., Themmen, A. P., 1983. Why are two different types of pyridine nucleotide transhydrogenase found in living organisms? Eur. J. Biochem. 131, 527–533.

Wilkes, R. A., Waldbauer, J., Carroll, A., Nieto-Domínguez, M., Parker, D. J., Zhang, L., Guss, A. M., Aristilde, L., 2023. Complex regulation in a *Comamonas* platform for diverse aromatic carbon metabolism. Nat. Chem. Biol. 19, 651–662.

Wirth, N. T., Kozaeva, E., Nikel, P. I., 2020. Accelerated genome engineering of *Pseudomonas putida* by I-*Sce*I―mediated recombination and CRISPR-Cas9 counterselection. Microb. Biotechnol. 13, 233–249.

Wiśniewski, J. R., Hein, M. Y., Cox, J., Mann, M., 2014. A "proteomic ruler" for protein copy number and concentration estimation without spike-in standards. Mol. Cell. Proteom. 13, 3497–3506.

Xu, Y., Guo, J., Jin, X., Kim, J. S., Ji, Y., Fan, S., Ha, N. C., Quan, C. S., 2016. Crystal structure and functional implications of the tandem-type universal stress protein UspE from *Escherichia coli*. BMC Struct. Biol. 16, 3.

Xuan, Y., Bateman, N. W., Gallien, S., Goetze, S., Zhou, Y., Navarro, P., Hu, M., Parikh, N., Hood, B. L., Conrads, K. A., Loosse, C., Kitata, R. B., Piersma, S. R., Chiasserini, D., Zhu, H., Hou, G., Tahir, M., Macklin, A., Khoo, A., Sun, X., Crossett, B., Sickmann, A., Chen, Y. J., Jimenez, C. R., Zhou, H., Liu, S., Larsen, M. R., Kislinger, T., Chen, Z., Parker, B. L., Cordwell, S. J., Wollscheid, B., Conrads, T. P., 2020. Standardization and harmonization of distributed multi-center proteotype analysis supporting precision medicine studies. Nat. Commun. 11, 5248.

Yishai, O., Lindner, S. N., González de la Cruz, J., Tenenboim, H., Bar-Even, A., 2016. The formate bio-economy. Curr. Opin. Chem. Biol. 35, 1–9.

Zhou, Y., Wang, L., Yang, F., Lin, X., Zhang, S., Zhao, Z. K., 2011. Determining the extremes of the cellular NAD(H) level by using an *Escherichia coli* NAD^+^-auxotrophic mutant. Appl. Environ. Microbiol. 77, 6133–6140.

Zhu, K., Zhang, Y. M., Rock, C. O., 2009. Transcriptional regulation of membrane lipid homeostasis in *Escherichia coli*. J. Biol. Chem. 284, 34880–34888.

Zobel, S., Kuepper, J., Ebert, B., Wierckx, N., Blank, L. M., 2017. Metabolic response of *Pseudomonas putida* to increased NADH regeneration rates. Eng. Life Sci. 17, 47–57.

