## Supplemental Data for "Integrated control of redox and energy metabolism by the membrane-bound and soluble transhydrogenases of *Pseudomonas putida* across metabolic regimes"

by

Michele Partipilo[1], Giusi Favoino[1], Òscar Puiggené[1], Catarina Rocha[1], Carina Meiners[2], Nicolás Gurdo[1], Stefano Donati[1], Daniel C. Volke[1], and Pablo I. Nikel[1*]

[1] The Novo Nordisk Foundation Center for Biosustainability, Technical University of Denmark, 2800 Kongens Lyngby, Denmark

[2] TUM School of Engineering and Design, Systems Biotechnology, Technical University of Munich, 85748 Garching, Germany

**Keywords:** transhydrogenases, redox reactions, nicotinamide adenine dinucleotides, metabolism, cell fitness, oxidative stress, *Pseudomonas putida*.

#### Supplementary Information

##### *Table of contents*

- Supplementary Tables S1-S8.
- Supplementary Figures S1-S9.
- Supplementary references.

#### Supplementary Tables

**Table S1.** Growth rates and maximal optical density values of wild-type *P. putida* and transhydrogenase mutants.

| Carbon source | Total γ | Growth rate ($h^{-1}$) | | | | Maximum OD | | | |
|---|---|---|---|---|---|---|---|---|---|
| | | WT | Δ*pntAB* | Δ*sthA* | Δ*sthA* Δ*pntAB* | WT | Δ*pntAB* | Δ*sthA* | Δ*sthA* Δ*pntAB* |
| Octanoate | 46.0 | 0.65 ± 0.09 | 0.68 ± 0.12 | 0.63 ± 0.15 | 0.26 ± 0.03 | 3.33 ± 0.04 | 3.45 ± 0.02 | 3.37 ± 0.14 | 1.63 ± 0.49 |
| Benzoate | 32.0 | 0.53 ± 0.01 | 0.57 ± 0.06 | 0.44 ± 0.01 | 0.10 ± 0.03 | 1.96 ± 0.04 | 1.95 ± 0.05 | 2.10 ± 0.07 | 0.22 ± 0.06 |
| Glucose | 24.0 | 0.73 ± 0.03 | 0.73 ± 0.08 | 0.76 ± 0.04 | 0.50 ± 0.03 | 2.80 ± 0.05 | 2.86 ± 0.03 | 2.84 ± 0.04 | 2.71 ± 0.07 |
| Fructose | 24.0 | 0.49 ± 0.04 | 0.58 ± 0.04 | 0.58 ± 0.01 | 0.29 ± 0.06 | 2.67 ± 0.06 | 2.62 ± 0.02 | 2.68 ± 0.01 | 2.45 ± 0.06 |
| Citrate | 18.0 | 0.87 ± 0.04 | 0.83 ± 0.07 | 0.85 ± 0.06 | 0.33 ± 0.03 | 2.15 ± 0.08 | 2.17 ± 0.09 | 2.23 ± 0.08 | 1.71 ± 0.18 |
| Glutamate | 18.0 | 0.86 ± 0.03 | 0.87 ± 0.02 | 0.84 ± 0.03 | 0.53 ± 0.06 | 2.57 ± 0.03 | 2.56 ± 0.03 | 2.57 ± 0.04 | 2.07 ± 0.19 |
| Ribose | 20.0 | 0.28 ± 0.07 | 0.35 ± 0.10 | 0.33 ± 0.12 | 0.19 ± 0.07 | 1.10 ± 0.03 | 1.12 ± 0.20 | 1.13 ± 0.08 | 1.15 ± 0.13 |
| Succinate | 14.0 | 0.65 ± 0.19 | 0.71 ± 0.20 | 0.70 ± 0.14 | 0.43 ± 0.11 | 2.40 ± 0.11 | 2.45 ± 0.11 | 2.48 ± 0.08 | 2.13 ± 0.15 |
| Lactate | 11.0 | 0.70 ± 0.01 | 0.77 ± 0.05 | 0.72 ± 0.02 | 0.44 ± 0.05 | 2.64 ± 0.06 | 2.61 ± 0.01 | 2.62 ± 0.04 | 2.15 ± 0.05 |
| Pyruvate | 9.0 | 0.53 ± 0.16 | 0.49 ± 0.17 | 0.40 ± 0.11 | 0.25 ± 0.04 | 2.21 ± 0.01 | 2.18 ± 0.02 | 2.05 ± 0.05 | 2.15 ± 0.05 |
| Glycerol | 14.0 | 0.52 ± 0.11 | 0.52 ± 0.10 | 0.58 ± 0.17 | 0.30 ± 0.08 | 2.59 ± 0.03 | 2.60 ± 0.09 | 2.61 ± 0.07 | 2.51 ± 0.19 |
| Acetate | 8.0 | 0.59 ± 0.03 | 0.53 ± 0.01 | 0.00 ± 0.00 | 0.02 ± 0.03 | 2.01 ± 0.03 | 2.02 ± 0.02 | 0.00 ± 0.00 | 0.01 ± 0.01 |

Cells were cultivated in minimal salt medium (MSM) at 30°C with different carbon sources, providing a total of 120 mM carbon atoms (corresponding to 15 mM octanoate, 17 mM benzoate, 20 mM glucose/fructose/citrate, 24 mM glutamate/ribose, 30 mM succinate, 40 mM lactate/pyruvate/glycerol, or 60 mM acetate) for 72 h, with shaking at 250 rpm. The growth parameters were obtained from biological replicates ($n$ = 4). Kinetic parameters, calculated employing the QurvE software (Wirth et al., 2023), are given as average values ± standard deviation. For each substrate, the degree of reduction (total γ) is shown, where total γ stands for the number of reducing equivalents available per carbon atom in the given molecule. Significant differences with the WT are highlighted in red as calculated by means of the Student's $t$ test with $p$-value ≤ 0.05.

**Table S2.** Cofactor ratios of the transhydrogenase strains on glucose and acetate.

| Cofactor ratio | On glucose | | | | On acetate | | | |
|---|---|---|---|---|---|---|---|---|
| | **WT** | **Δ*pntAB*** | **Δ*sthA*** | **Δ*sthA* Δ*pntAB*** | **WT** | **Δ*pntAB*** | **Δ*sthA*** | **Δ*sthA* Δ*pntAB*** |
| $NADH/NAD^+$ | 0.008 ± 0.001 | 0.008 ± 0.000 | 0.008 ± 0.001 | 0.007 ± 0.005 | 0.031 ± 0.006 | 0.026 ± 0.000 | N.D. | N.D. |
| $NADPH/NADP^+$ | 0.43 ± 0.01 | 0.40 ± 0.06 | 0.44 ± 0.04 | 0.31 ± 0.07 | 1.41 ± 0.45 | 1.70 ± 0.24 | N.D. | N.D. |
| ATP/ADP | 15.6 ± 4.5 | 13.7 ± 2.4 | 16.8 ± 1.9 | 8.1 ± 1.9 | 5.3 ± 2.3 | 5.1 ± 0.3 | N.D. | N.D. |
| Energy charge | 0.93 ± 0.01 | 0.93 ± 0.01 | 0.94 ± 0.01 | 0.87 ± 0.03 | 0.77 ± 0.11 | 0.83 ± 0.01 | N.D. | N.D. |

Cells samples were grown in MSM at 30°C with either 20 mM glucose or 60 mM acetate as the sole carbon source and then treated for absolute quantification on LC-MS. Measurements come from biological triplicates, each of which performed in technical duplicate, while errors represent standard deviation. N.D., not detected.

**Table S3.** List of PntAB and SthA sequences used for phylogenetic analysis. The taxonomical origin is reported for each representative chosen organism. Shades indicate the taxonomical order (Pseudomonadales in yellow, Enterobacterales in light blue, Vibrionales in light brown, Hyphomicrobiales in light green, Mycobacteriales in light red, Alteromonadales in white).

| Organism | Family | Order | Class | Phylum | UniProt SthA | UniProt PntB |
|---|---|---|---|---|---|---|
| *Acinetobacter baumanii* | Moraxellaceae | Pseudomonadales | Gammaproteobacteria | Pseudomonadota | V5VCW4 | V5VI87 |
| *Acinetobacter haemolyticus* | Moraxellaceae | Pseudomonadales | Gammaproteobacteria | Pseudomonadota | D4XPK1 | D4XMM7 |
| *Acinetobacter lwoffii* | Moraxellaceae | Pseudomonadales | Gammaproteobacteria | Pseudomonadota | A0A2K8UK16 | A0AAJ4P4S2 |
| *Agrobacterium vitis* | Rhizobiaceae | Hyphomicrobiales | Alphaproteobacteria | Pseudomonadota | A0AAN1X2R7 | A0AAE4WL10 |
| *Aliivibrio fischeri ES114* | Vibrionaceae | Vibrionales | Gammaproteobacteria | Pseudomonadota | Q5E212 | Q5DZZ1 |
| *Alteromonas macleodii* | Alteromonadaceae | Alteromonadales | Gammaproteobacteria | Pseudomonadota | A0A6T9Y3H7 | A0A6T9Y7F5 |
| *Azotobacter vinelandii* | Pseudomonadaceae | Pseudomonadales | Gammaproteobacteria | Pseudomonadota | C1DR10 | C1DH39 |
| *Citrobacter amalonaticus* | Enterobacteriaceae | Enterobacterales | Gammaproteobacteria | Pseudomonadota | A0A0F6TYE6 | M1K5K1 |
| *Citrobacter freundii* | Enterobacteriaceae | Enterobacterales | Gammaproteobacteria | Pseudomonadota | A0A7D6XWK2 | A0A7D6ZRD1 |
| *Citrobacter koseri ATCC BAA-895* | Enterobacteriaceae | Enterobacterales | Gammaproteobacteria | Pseudomonadota | A8AKW0 | A8AGX9 |
| *Cronobacter sakazakii* | Enterobacteriaceae | Enterobacterales | Gammaproteobacteria | Pseudomonadota | A7ML92 | A7MMP5 |
| *Cronobacter turicensis* | Enterobacteriaceae | Enterobacterales | Gammaproteobacteria | Pseudomonadota | C9Y4R2 | C9Y3Q9 |
| *Dickeya dadantii* | Pectobacteriaceae | Enterobacterales | Gammaproteobacteria | Pseudomonadota | E0SGD3 | E0SCK6 |
| *Edwardsiella piscicida* | Hafniaceae | Enterobacterales | Gammaproteobacteria | Pseudomonadota | A0AAU8P706 | A0AAU8PC33 |
| *Edwardsiella tarda* | Hafniaceae | Enterobacterales | Gammaproteobacteria | Pseudomonadota | A0A0H3DXG7 | A0A0H3DTE3 |
| *Ensifer adhaerens* | Rhizobiaceae | Hyphomicrobiales | Alphaproteobacteria | Pseudomonadota | A0A9Q8YB94 | A0A9Q8Y7R4 |
| *Enterobacter cloacae* | Enterobacteriaceae | Enterobacterales | Gammaproteobacteria | Pseudomonadota | A0A427KKJ5 | Q19KD3 |
| *Enterobacter lignolyticus* | Enterobacteriaceae | Enterobacterales | Gammaproteobacteria | Pseudomonadota | E3G430 | E3G6S6 |
| *Erwinia persicina* | Erwiniaceae | Enterobacterales | Gammaproteobacteria | Pseudomonadota | A0A3S7S8D1 | A0A354AEC0 |
| *Enterobacter soli* | Enterobacteriaceae | Enterobacterales | Gammaproteobacteria | Pseudomonadota | A0AAW8HDM3 | A0AAW8HAG6 |
| *Escherichia coli str. K-12 substr. MG1655* | Enterobacteriaceae | Enterobacterales | Gammaproteobacteria | Pseudomonadota | P27306 | P0AB67 |
| *Hafnia alvei* | Hafniaceae | Enterobacterales | Gammaproteobacteria | Pseudomonadota | G9Y9A0 | G9Y1N8 |
| *Kluyvera ascorbata* | Enterobacteriaceae | Enterobacterales | Gammaproteobacteria | Pseudomonadota | A0A378GES6 | A0A378GR38 |

| | | | | | | |
|---|---|---|---|---|---|---|
| *Lelliottia amnigena* | Enterobacteriaceae | Enterobacterales | Gammaproteobacteria | Pseudomonadota | A0AAP2F4B7 | A0AAP2EZ40 |
| *Marinobacter nauticus ATCC 49840* | Alteromonadaceae | Alteromonadales | Gammaproteobacteria | Pseudomonadota | A1U1Y5 | A1TXK3 |
| *Mycobacterium marinum E11* | Mycobacteriaceae | Mycobacteriales | Actinomycetes | Actinomycetota | B2HLY6 | B2HLU7 |
| *Mycobacterium tuberculosis H37Rv* | Mycobacteriaceae | Mycobacteriales | Actinomycetes | Actinomycetota | A5U665 | A5TYN2 |
| *Mycobacteroides chelonae CCUG 47445* | Mycobacteriaceae | Mycobacteriales | Actinomycetes | Actinomycetota | A0AA40XFQ2 | A0A1S1M8Y0 |
| *Mycolicibacterium fortuitum subsp. fortuitum* | Mycobacteriaceae | Mycobacteriales | Actinomycetes | Actinomycetota | A0AAW4KGJ5 | A0AAW4KD45 |
| *Obesumbacterium proteus* | Hafniaceae | Enterobacterales | Gammaproteobacteria | Pseudomonadota | A0AA91EDC2 | A0AA91EC24 |
| *Pantoea (Enterobacter) agglomerans* | Erwiniaceae | Enterobacterales | Gammaproteobacteria | Pseudomonadota | A0A6I6K4F1 | A0AAN2FBR3 |
| *Pantoea stewartii* | Erwiniaceae | Enterobacterales | Gammaproteobacteria | Pseudomonadota | H3RI37 | H3RD07 |
| *Pectobacterium atrosepticum* | Pectobacteriaceae | Enterobacterales | Gammaproteobacteria | Pseudomonadota | Q6CZB1 | Q6D537 |
| *Pectobacterium carotovorum* | Pectobacteriaceae | Enterobacterales | Gammaproteobacteria | Pseudomonadota | A0AA40J1B6 | A0AA40MA20 |
| *Photobacterium damselae subsp. damselae* | Vibrionaceae | Vibrionales | Gammaproteobacteria | Pseudomonadota | D0Z0P8 | D0Z2G5 |
| *Photobacterium phosphoreum* | Vibrionaceae | Vibrionales | Gammaproteobacteria | Pseudomonadota | A0A2T3JJE4 | A0A2T3PIZ9 |
| *Photorhabdus laumondii subsp. laumondii* | Morganellaceae | Enterobacterales | Gammaproteobacteria | Pseudomonadota | Q7MBG9 | Q7N4Z2 |
| *Proteus mirabilis HI4320* | Morganellaceae | Enterobacterales | Gammaproteobacteria | Pseudomonadota | B4F1H1 | B4ETT1 |
| *Proteus vulgaris* | Morganellaceae | Enterobacterales | Gammaproteobacteria | Pseudomonadota | A0A6G6SD87 | A0A6G6SHJ5 |
| *Providencia rettgeri* | Morganellaceae | Enterobacterales | Gammaproteobacteria | Pseudomonadota | A0A1B8SWG1 | A0A379FRR4 |
| *Providencia stuartii* | Morganellaceae | Enterobacterales | Gammaproteobacteria | Pseudomonadota | A0AA86YQS8 | A0AA86YJF6 |
| *Pseudomonas aeruginosa* | Pseudomonadaceae | Pseudomonadales | Gammaproteobacteria | Pseudomonadota | P57112 | Q9I6T9 |
| *Pseudomonas putida* | Pseudomonadaceae | Pseudomonadales | Gammaproteobacteria | Pseudomonadota | Q88KY8 | Q88RH4 |
| *Pseudomonas fluorescens* | Pseudomonadaceae | Pseudomonadales | Gammaproteobacteria | Pseudomonadota | C3K4W1 | C3K7Y9 |
| *Pseudomonas syringae* | Pseudomonadaceae | Pseudomonadales | Gammaproteobacteria | Pseudomonadota | Q884I6 | Q87U51 |
| *Raoultella ornithinolytica* | Enterobacteriaceae | Enterobacterales | Gammaproteobacteria | Pseudomonadota | A0A1Y6GPH2 | A0A1Y6GGK7 |
| *Rhizobium leguminosarum* | Rhizobiaceae | Hyphomicrobiales | Alphaproteobacteria | Pseudomonadota | A0A2K9Z1L9 | A0A2K9Z898 |
| *Salmonella typhi* | Enterobacteriaceae | Enterobacterales | Gammaproteobacteria | Pseudomonadota | P66009 | Q8XGU1 |
| *Serratia liquefaciens ATCC 27592* | Yersiniaceae | Enterobacterales | Gammaproteobacteria | Pseudomonadota | A0A380AVA5 | A0A515CUU5 |
| *Serratia marcescens* | Yersiniaceae | Enterobacterales | Gammaproteobacteria | Pseudomonadota | A0AAT9F551 | A0AAT9EMU9 |

| | | | | | | |
|---|---|---|---|---|---|---|
| *Shigella dysenteriae* | Enterobacteriaceae | Enterobacterales | Gammaproteobacteria | Pseudomonadota | Q32AB3 | Q32G68 |
| *Stutzerimonas stutzeri* | Pseudomonadaceae | Pseudomonadales | Gammaproteobacteria | Pseudomonadota | A4VMU6 | A4VFS8 |
| *Vibrio cholerae* | Vibrionaceae | Vibrionales | Gammaproteobacteria | Pseudomonadota | P50529 | Q9KM25 |
| *Vibrio coralliilyticus* | Vibrionaceae | Vibrionales | Gammaproteobacteria | Pseudomonadota | A0AAN0S9G0 | A0AAN0SHG9 |
| *Vibrio natriegens NBRC 15636 = ATCC 14048 = DSM 759* | Vibrionaceae | Vibrionales | Gammaproteobacteria | Pseudomonadota | A0AAN0Y4R0 | A0AAN0Y615 |
| *Vibrio vulnificus NBRC 15645 = ATCC 27562* | Vibrionaceae | Vibrionales | Gammaproteobacteria | Pseudomonadota | Q7MQ83 | Q7ME73 |
| *Xenorhabdus bovienii subsp. africana* | Morganellaceae | Enterobacterales | Gammaproteobacteria | Pseudomonadota | A0A0B6XGK5 | A0A0B6X9V8 |
| *Yersinia enterocolitica subsp. palearctica Y11* | Yersiniaceae | Enterobacterales | Gammaproteobacteria | Pseudomonadota | A1JI37 | A1JN46 |
| *Yersinia pestis A1122* | Yersiniaceae | Enterobacterales | Gammaproteobacteria | Pseudomonadota | Q8ZA97 | A0A0H2W604 |
| *Yersinia ruckeri* | Yersiniaceae | Enterobacterales | Gammaproteobacteria | Pseudomonadota | A0A085U3J3 | A0A085U7N0 |

**Table S4.** Genomic context of the membrane-bound and soluble transhydrogenases from selected representative organisms sharing the dual transhydrogenase adaptation. For each strain, we retrieved the location of both *pntB* and *sthA* on https://pubseed.theseed.org/FIG/seedviewer.cgi?page=Minimal and consulted the annotation of the neighboring genes, reported on **Fig. 6D** of the main text.

| No. | Strain | *pntB* | *sthA* |
|---|---|---|---|
| 1 | *Pseudomonas putida* KT2440 | https://pubseed.theseed.org/FIG/seedviewer.cgi?page=Annotation&feature=fig%7C160488.1.peg.155 | https://pubseed.theseed.org/FIG/seedviewer.cgi?page=Annotation&feature=fig%7C160488.1.peg.2130 |
| 2 | *Pseudomonas aeruginosa* PAO1 | https://pubseed.theseed.org/FIG/seedviewer.cgi?page=Annotation&feature=fig\|208964.1.peg.197 | https://pubseed.theseed.org/FIG/seedviewer.cgi?page=Annotation&feature=fig\|208964.1.peg.2991 |
| 3 | *Acinetobacter baumannii* ATCC 17978 | https://pubseed.theseed.org/FIG/seedviewer.cgi?page=Annotation&feature=fig\|400667.4.peg.567 | https://pubseed.theseed.org/FIG/seedviewer.cgi?page=Annotation&feature=fig\|400667.4.peg.2437 |
| 4 | *Mycobacterium tuberculosis* | https://pubseed.theseed.org/FIG/seedviewer.cgi?page=Annotation&feature=fig\|1773.31.peg.2468 | https://pubseed.theseed.org/FIG/seedviewer.cgi?page=Annotation&feature=fig\|1773.31.peg.267 |
| 5 | *Rhizobium leguminosarum* bv. viciae 3841 | https://pubseed.theseed.org/FIG/seedviewer.cgi?page=Annotation&feature=fig\|216596.1.peg.4269 | https://pubseed.theseed.org/FIG/seedviewer.cgi?page=Annotation&feature=fig\|216596.1.peg.2153 |
| 6 | *Alteromonas macleodii* 'Deep ecotype' | https://pubseed.theseed.org/FIG/seedviewer.cgi?page=Annotation&feature=fig\|314275.3.peg.3378 | https://pubseed.theseed.org/FIG/seedviewer.cgi?page=Annotation&feature=fig\|314275.3.peg.1984 |
| 7 | *Vibrio cholerae* O1 biovar eltor str. N16961 | https://pubseed.theseed.org/FIG/seedviewer.cgi?page=Annotation&feature=fig\|243277.26.peg.3191 | https://pubseed.theseed.org/FIG/seedviewer.cgi?page=Annotation&feature=fig%7C243277.1.peg.150 |
| 8 | *Photorhabdus luminescens* subsp. *laumondii* TTO1 | https://pubseed.theseed.org/FIG/seedviewer.cgi?page=Annotation&feature=fig\|243265.1.peg.2052 | https://pubseed.theseed.org/FIG/seedviewer.cgi?page=Annotation&feature=fig\|243265.1.peg.4524 |
| 9 | *Yersinia pestis* CO92 | https://pubseed.theseed.org/FIG/seedviewer.cgi?page=Annotation&feature=fig\|214092.1.peg.2355 | https://pubseed.theseed.org/FIG/seedviewer.cgi?page=Annotation&feature=fig%7C214092.1.peg.3875 |
| 10 | *Dickeya dadantii* 3937 | https://pubseed.theseed.org/FIG/seedviewer.cgi?page=Annotation&feature=fig\|198628.6.peg.2309 | https://pubseed.theseed.org/FIG/seedviewer.cgi?page=Annotation&feature=fig\|198628.6.peg.214 |
| 11 | *Enterobacter cloacae* SCF1 | https://pubseed.theseed.org/FIG/seedviewer.cgi?page=Annotation&feature=fig\|701347.4.peg.2281 | https://pubseed.theseed.org/FIG/seedviewer.cgi?page=Annotation&feature=fig\|701347.4.peg.4358 |
| 12 | *Escherichia coli* str. K-12 substr. MG1655 | https://pubseed.theseed.org/FIG/seedviewer.cgi?page=Annotation&feature=fig%7C511145.6.peg.1657 | https://pubseed.theseed.org/FIG/seedviewer.cgi?page=Annotation&feature=fig%7C511145.6.peg.4065 |

**Table S5.** Functional annotation of the genes found in the genomic contexts of selected microbes. For all organisms represented in Figure 6D of the main text, we annotated and categorized the function of both *sthA* and *pntAB* flanking genes. Each physiological function (carbon and amino acid metabolism, lipid homeostasis, redox metabolism, cell division, cell signaling, molecular transport, gene expression, cell stress, oxidative stress, respiration, extracellular electron, toxin-antitoxin system, not annotated function) is shaded in a specific color.

| Transhydrogenase gene | Neighboring gene | Class |
|---|---|---|
| *sthA* | 1: NADPH-dependent glyceraldehyde-3-phosphate dehydrogenase | Carbon metabolism |
| | 2: FAD:protein FMN transferase | Extracellular electron transfer |
| | 3: Glycerophosphoryl diester phosphodiesterase | Lipid homeostasis |
| | 4: PilZ domain protein, c-di-GMP-binding | Cell signalling |
| | 5: Na(+) translocating NADH quinone reductase subunit | Extracellular electron transfer |
| | 6: Probable exported or periplasmic protein in ApbE locus | Extracellular electron transfer |
| | 7: Phosphodiesterase | Cell signalling |
| | 8: Short-chain acyl-CoA dehydrogenase | Lipid homeostasis |
| | 9: Lipoyl synthase | Lipid homeostasis |
| | 10: Permeases of the major facilitator superfamily | Molecular transport |
| | 11: Nitrate/ nitrite transporter | Molecular transport |
| | 12: Iron dependent repressor IdeR/DtxR | Oxidative stress |
| | 13: DUF4192 domain-containing protein | Not annotated |
| | 14: Proteasom assembly chaperon family protein MSMEG_2746 | Cell stress |
| | 15: Hydrolase, alpha beta/fold family | Redox metabolism |
| | 16: DNA repair protein RadC | Cell stress |
| | 17: L-lactate dehydrogenase | Carbon metabolism |
| | 18: Endonuclease/exonuclease/phosphatase family metal-dependent hydrolase | Redox metabolism |
| | 19: P-methylase | Not annotated |
| | 20: Unsaturated fatty acid biosynthesis repressor FabR, TetR family | Lipid homeostasis |
| | 21: Fatty acid desaturase | Lipid homeostasis |
| | 22: Not annotated | Not annotated |
| | 23: Uracil-DNA glycosylase family protein | Cell stress |
| | 24: Inner membrane protein YijD | Lipid homeostasis |
| | 25: RNA polymerase sigma factor RpoH | Cell stress |
| | 26: Cell division associated, ABC transporter like signaling protein FtsX | Cell division |
| | 27: Hydrogen peroxide-inducible genes activator OxyR | Oxidative stress |

| | | |
|---|---|---|
| | 28: Argininosuccinate lyase | Carbon metabolism |
| | 29: Hybrid peroxiredoxin hyPrx5 | Oxidative stress |
| | 30: Peroxidredoxin family protein/ glutaredoxin | Oxidative stress |
| *pntAB* | 31: Glutaryl-CoA dehydrogenase | Carbon metabolism |
| | 32: Transcriptional regulator PA0448, LysR family | Gene expression |
| | 33: TPP riboswitch (THI element) | Gene expression |
| | 34: Acetyl-CoA hydrolase/transferase family protein | Carbon metabolism |
| | 35: Aliphatic sulfate esters (C4-C12 chain lengths) dioxygenase | Redox metabolism |
| | 36: Dioxygenase, TauD/ TfdA family | Redox metabolism |
| | 37: Ton complex, TonB subunit | Cell signalling |
| | 38: Ton complex ExbB subunit | Cell signalling |
| | 39: MFS permease protein | Molecular transport |
| | 40: Permease of the drug/ metabolite transporter (DMT) superfamily | Molecular transport |
| | 41: Retinol dehydrogenase | Redox metabolism |
| | 42: Ribosomal silencing factor RsfA | Gene expression |
| | 43: Acyl-CoA dehydrogenase, Mycobacterial subgroup FadE2 | Carbon metabolism |
| | 44: DUF4440 domain-containing protein | Not annotated |
| | 45: Possible outer membrane protein | Not annotated |
| | 46: Transcriptional regulator, AcrR family | Gene expression |
| | 47: Hypothetical protein | Not annotated |
| | 48: Aa3-type cytochrome c oxidase subunit IV | Respiration |
| | 49: DUF992 domain-containing protein | Not annotated |
| | 50: Sensor histidine kinase, putative | Cell signalling |
| | 51: Tryptophan 7 halogenase | Redox metabolism |
| | 52: Predicted b-glucoside-specific TonB-dependent outer membrane receptor | Cell signalling |
| | 53: hypothetical protein | Not annotated |
| | 54: Biosynthetic arginine decarboxylase | Carbon metabolism |
| | 55: Late competence protein ComFB | Cell signalling |
| | 56: Transcriptional activator, HlyU family | Gene expression |
| | 57: component sensor histidine kinase VCA0565 | Cell signalling |
| | 58: Two component transcriptional response regulator VCA0566, OmpR family | Cell signalling |
| | 59: Acyl carrier protein | Lipid homeostasis |
| | 60: Protein YdgH @ uncharacterized BhsA-like protein | Cell stress |
| | 61: Death on curing protein, Doc toxin | Toxin-Antitoxin system |

| | | |
|---|---|---|
| | 62: Prevent host death protein, PhD antitoxin | Toxin-Antitoxin system |
| | 63: Uncharacterized protein | Not annotated |
| | 64: Universal stress protein UspE | Cell stress |
| | 65: Fumarate and nitrate reduction regulatory protein Fnr | Oxidative stress |
| | 66: 2-keto 3-deoxy-D-manno-octulosonate-8-phosphate synthase | Lipid homeostasis |
| | 67: Arginine/ omithine antiporter ArcD | Molecular transport |
| | 68: Leucine efflux protein | Molecular transport |
| | 69: S-(hydroxymethyl) glutathione dehydrogenase | Oxidative stress |
| | 70: AI-2 transport protein TqsA | Molecular transport |
| | 71: Spermidine export protein MdtJ | Molecular transport |

**Table S6.** Bacterial strains used in this study.

| Strain | Relevant characteristics[a] | Reference or source |
|---|---|---|
| *Escherichia coli* | | |
| DH5α λ*pir* | Cloning host; F⁻ λ⁻ *endA1 glnX44(AS) thiE1 recA1 relA1 spoT1* *gyrA96*(Nal$^R$) *rfbC1 deoR nupG* F*80(lacZ*Δ*M15)* Δ*(argF-lac)U169* *hsdR17(*$r_K^-$ $m_K^+$*)*, λ*pir* lysogen | (Platt et al., 2000) |
| HB101 | Helper strain; F⁻ *thi*-1 *hsd*S20 ($r_B$ – $m_B$ – ) *sup*E44 *rec*A13 *ara*-14 *leu*B6 *pro*A2 *lac*Y1 *galK*2 *rpsL*20(Str$^R$) *xyl*-5 *mtl*-1 | (Kessler et al., 1992) |
| *Pseudomonas putida* | | |
| KT2440 | Parental strain, derived from *P. putida* mt-2 cured of the TOL plasmid pWW0 | (Bagdasarian et al., 1981) |
| Δ*pntAB* | Derivative of *P. putida* KT2440, Δ*pntAB* | (Nikel et al., 2016) |
| Δ*sthA* | Derivative of *P. putida* KT2440, Δ*sthA* | (Nikel et al., 2016) |
| Δ*pntAB* Δ*sthA* | Derivative of *P. putida* KT2440, Δ*pntAB* Δ*sthA* | (Nikel et al., 2016) |
| Δ*pntAB* Δ*PP_0256-0257* | Derivative of *P. putida* KT2440, Δ*pntAB* Δ*PP_0256-57* | This work |
| Δ*sthA* Δ*PP_0256-0257* | Derivative of *P. putida* KT2440, Δ*sthA* Δ*PP_0256-57* | This work |
| Δ*pntAB* Δ*sthA* Δ*PP_0256-0257* | Derivative of *P. putida* KT2440, Δ*pntAB* Δ*sthA* Δ*PP_0256-57* | This work |
| Δ*sthA* A1C1 (ALE #1 clone 1) | Derivative of *P. putida* KT2440, Δ*sthA* Δ*PP_1261-1264* | This work |
| Δ*sthA* A2C1 (ALE #2 clone 1) | Derivative of *P. putida* KT2440, Δ*sthA* Δ*PP_1261-1262* | This work |
| Δ*sthA* A2C2 (ALE #2 clone 2) | Derivative of *P. putida* KT2440, Δ*sthA* Δ*PP_1261* with truncated *PP_1262* (missing region 1,442,061 - 1,443,672 bp – only the first 171/315 amino acids of PP_1262 can be translated) | This work |
| Δ*sthA* A2C3 (ALE #2 clone 3) | Derivative of *P. putida* KT2440, Δ*sthA* with premature stop codon in *PP_1262* (stop codon at position 1,444,053 bp – CAA codon becomes TAA, resulting in the truncated PP_1262 Q45*) | This work |

[a]Antibiotic markers: Nal, nalidixic acid; Str, streptomycin.

**Table S7.** Plasmids used in this study.

| Plasmid | Relevant characteristics[a] | Reference or source |
|---|---|---|
| pSNW2 | Suicide vector used for deletions in Gram-negative bacteria; *oriT*, *traJ*, *lacZ*α, *ori*(R6K), $P_{14g}$(BCD2)→*msfGFP*; $Km^R$ | (Volke et al., 2020) |
| pSNW2_Δ*PP_1261* | Derivative of pSNW2 for deleting *PP_1261*; $Km^R$ | This work |
| pGNW2_Δ*PP_0256-57* | Derivative of pGNW2 for deleting *PP_0256-57*; $Km^R$ | (Turlin et al., 2023) |
| pQURE6·H | Helper plasmid for gene deletions; conditionally replicating vector carrying XylS/*Pm*→*I-SceI* and $P_{14g}$(BCD2)→*mRFP*; $Gm^R$ | (Volke et al., 2020) |
| pS621c | Control vector, carrying a $P_{trc}$ promoter and the canonical SEVA ribosome binding site (RBS); *oriT, oriV*(RK2); $Gm^R$ | (Turlin et al., 2023) |
| pFDH1 | Derivative of pS621c, $P_{trc}$ →carrying a codon optimized *Snov3272* from *Ancylobacter novellus* DSM 506 with 10X-his tag at the C-terminus ;(Partipilo et al., 2021) *oriT oriV*(RK2); $Gm^R$ | This work |
| pFDH2 | Derivative of pS621c, $P_{trc}$ →codon optimized *Snov3272* from *Ancylobacter novellus* DSM 506 with 10X-his tag at the C-terminus with mutations A198G/D221Q/H379K/S380V in the open reading frame;(Partipilo et al., 2023) *oriT oriV*(RK2); $Gm^R$ | This work |

[a]Antibiotic markers: Gm, gentamicin; Km, kanamycin

**Table S8.** Primers used in this study.

| Name | DNA sequence (5'→3') | Use |
|---|---|---|
| Primer_001 | AACATAUGAGTGCCAAAATTCTGTG | Amplification of of $NAD^+$-Fdh and $NADP^+$-Fdh for cloning into pS621c |
| Primer_002 | ACATTTAAUGATGATGATGATGGTGATG | |
| Primer_003 | ATTAAATGUAATCTAGAGTCGACCTGC | Amplification of pS621c for cloning with Fdh-variants |
| Primer_004 | ATATGTUTTTCCTCCTGGGAATTCG | |
| Primer_005 | AGCGGATAACAATTTCACACAGGA | Sequencing of inserts into pS621 plasmids |
| Primer_006 | GTCGTGACTGGGAAAACCCTGGCG | |
| Primer_007 | GCTGAAGGCGGTTGGTCGCAAG | Verification of Δ*sthA* in *P. putida* KT2440 |
| Primer_008 | CGACAACGAACTGATGGTCATCCACGAC | |
| Primer_009 | GGCTGGTGGTCGCTAACAGGCATG | Verification of Δ*pntAB* in *P. putida* KT2440 |
| Primer_010 | GGTTGGTGTTGTGCGAGAGGTTGATCTC | |
| Primer_011 | AGATCCUAGAGATGGTCAACATCACCAGTT | Construction of pSNW2·ΔPP_1261 |
| Primer_012 | AGCCCTTCUTTTTCATCGGGCAACTTCTCT | |
| Primer_013 | AGAAGGGCUGACCGTTTGATGGTGCCTTTG | |
| Primer_014 | AGG TCG ACU CAGGTCCGGGCTTTGCTT | |
| Primer_015 | GGAGAAGTGGCTGTTCGATGGC | Verification of Δ*PP_1261* |
| Primer_016 | GGAGTTCGAAACCAGAAGCCAAGAC | |
| Primer_017 | GGCATTGCCATACACGACCATGG | Verification of Δ*PP_0256-257* |
| Primer_018 | CTCTTTGCCCAGGTAGTTACCCAGCG | |

### Supplementary Figures

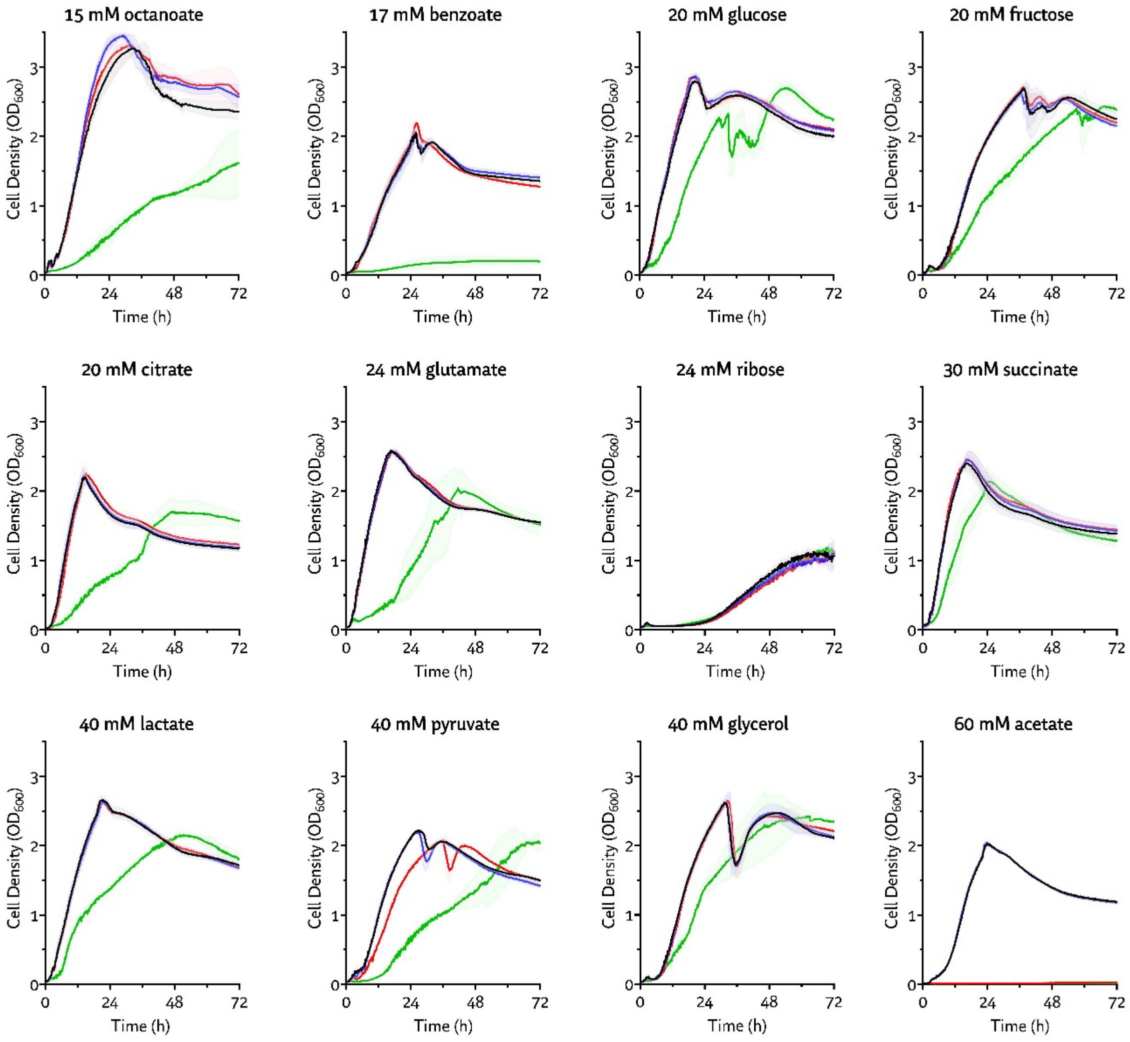


**Figure S1.** Growth curves of *P. putida* KT2440 WT and transhydrogenase mutant strains on different carbon sources. The color code is the same used in the main text (WT in black, Δ*pntAB* in blue, Δ*sthA* in red, Δ*pntAB* Δ*sthA* in green). The experiments were performed in biological quadruplicate (*n* = 4, errors represented as standard deviation).

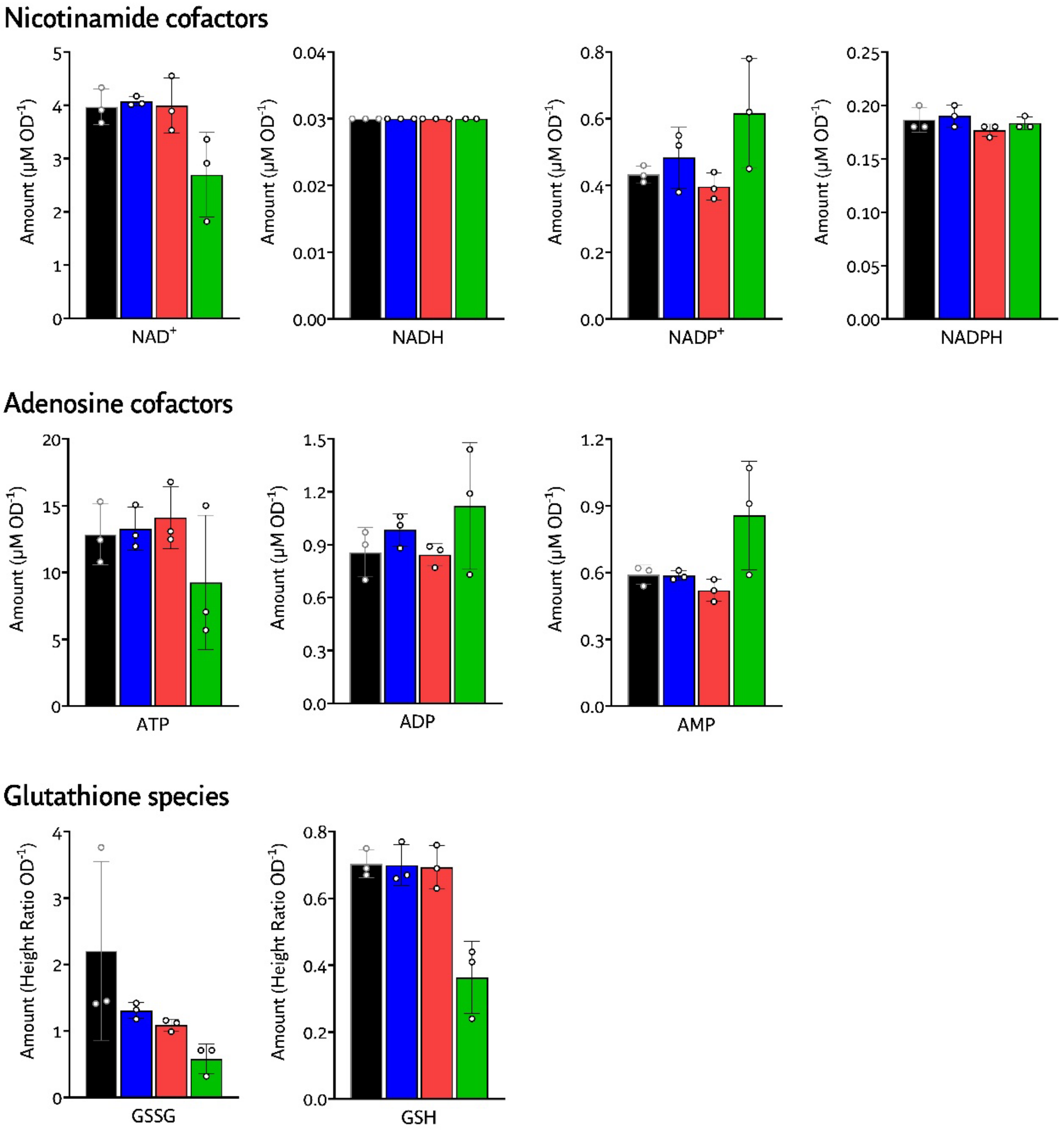


**Figure S2.** Measured amount of nicotinamide cofactors, adenosine cofactors and glutathione species on glucose. Cells were grown in minimal medium with 20 mM glucose as the sole carbon source and harvested for Liquid Chromatography-Mass Spectrometry analyses (see 'Material and Methods' section for details). In the case of $NAD^+$, NADH, $NADP^+$, NADPH, ATP, ADP and AMP the quantification was carried out on known standards, while the height ratio was used to estimate the amount of GSSG and GSH. Data were obtained in biological triplicate ($n$ = 3, errors shown as standard deviation). The color code is the same used in the main text (WT in black, Δ*pntAB* in blue, Δ*sthA* in red, Δ*pntAB* Δ*sthA* in green).

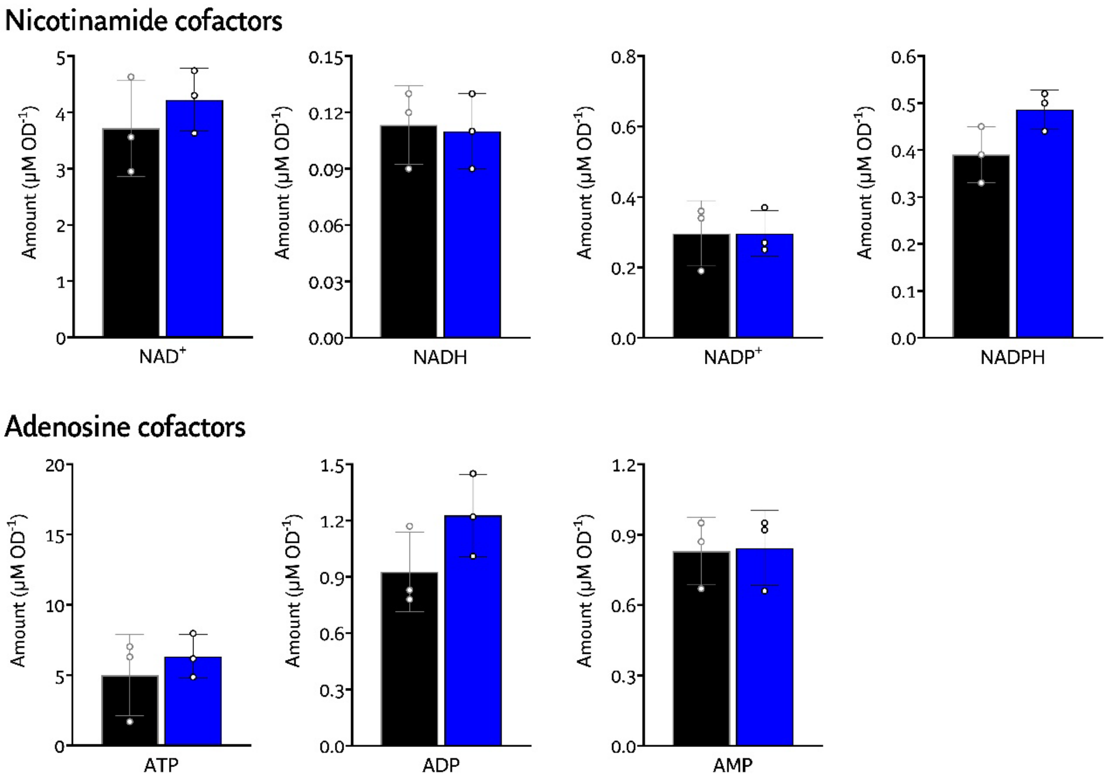


**Figure S3.** Measured amount of nicotinamide cofactors and adenosine cofactors on acetate. Cells were grown in minimal medium with 60 mM acetate as the sole carbon source and harvested for LC-MS analyses (see 'Material and Methods' section for details). Cofactors were quantified by known standards. Data were obtained in biological triplicate (*n* = 3, errors shown as standard deviation). The color code is the same used in the main text (WT in black and Δ*pntAB* in blue).

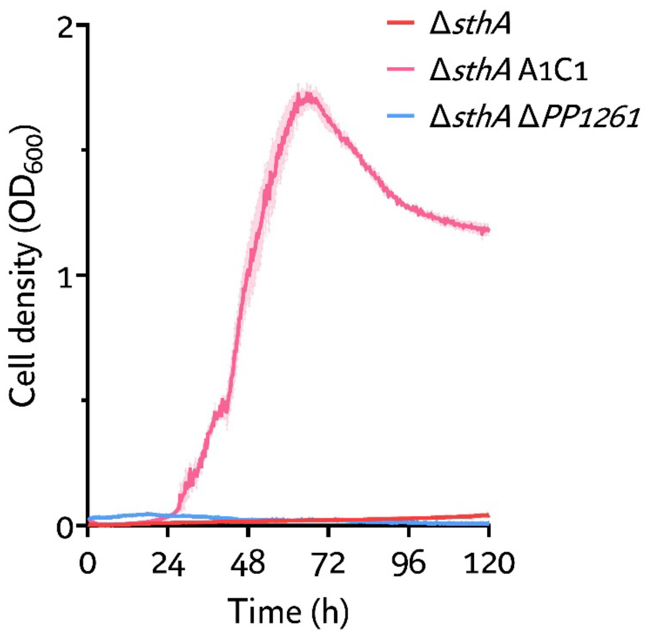


**Figure S4.** Growth curves of *P. putida* KT2440 Δ*sthA*, Δ*sthA* Δ*PP_1261* and ALE-derived Δ*sthA* A1C1 on 60 mM acetate. The experiments were performed in biological triplicate (*n* = 3, errors shown as standard deviation).

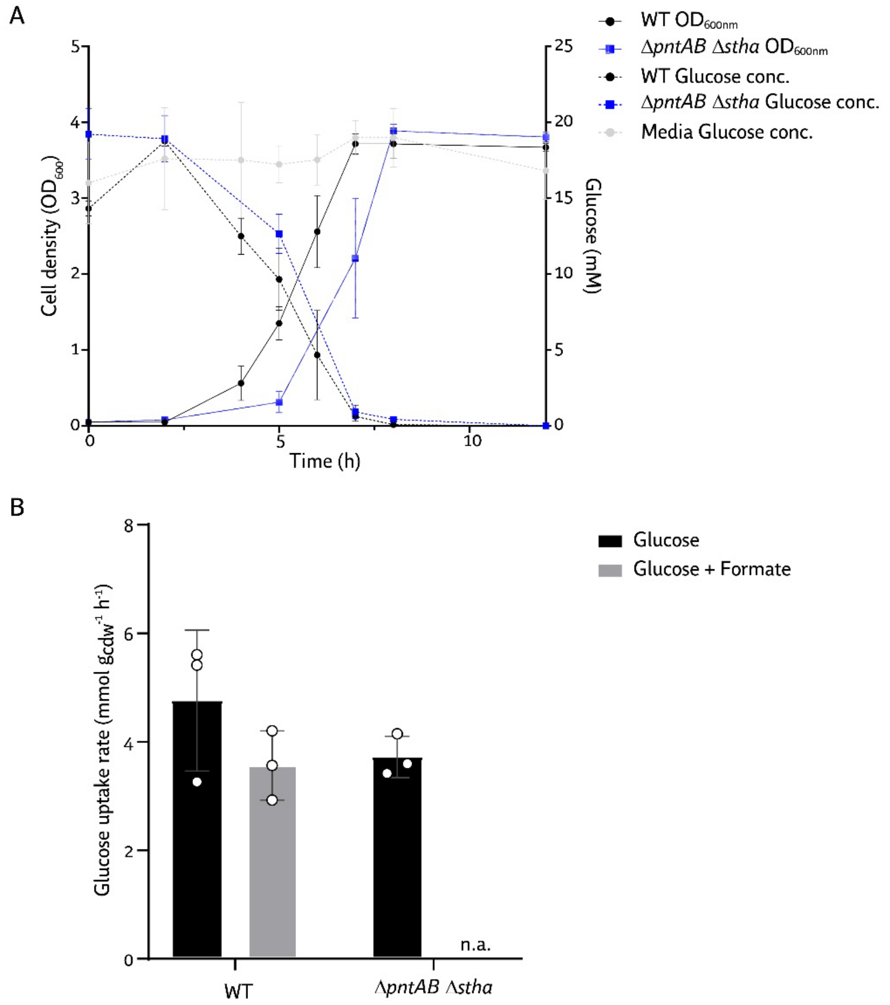


**Figure S5. A** Growth curves and glucose consumption of *P. putida* KT2440 WT and Δ*pntAB* Δ*sthA*. In MSM with 20 mM glucose. **B** Glucose uptake rate of WT and Δ*pntAB* Δ*sthA* in presence or absence of formate. The rate for the double mutant in glucose alone was calculated considering all the $OD_{600}$ points, and not only the exponential phase, due to missing data in the exponential phase. N.a.: not available.

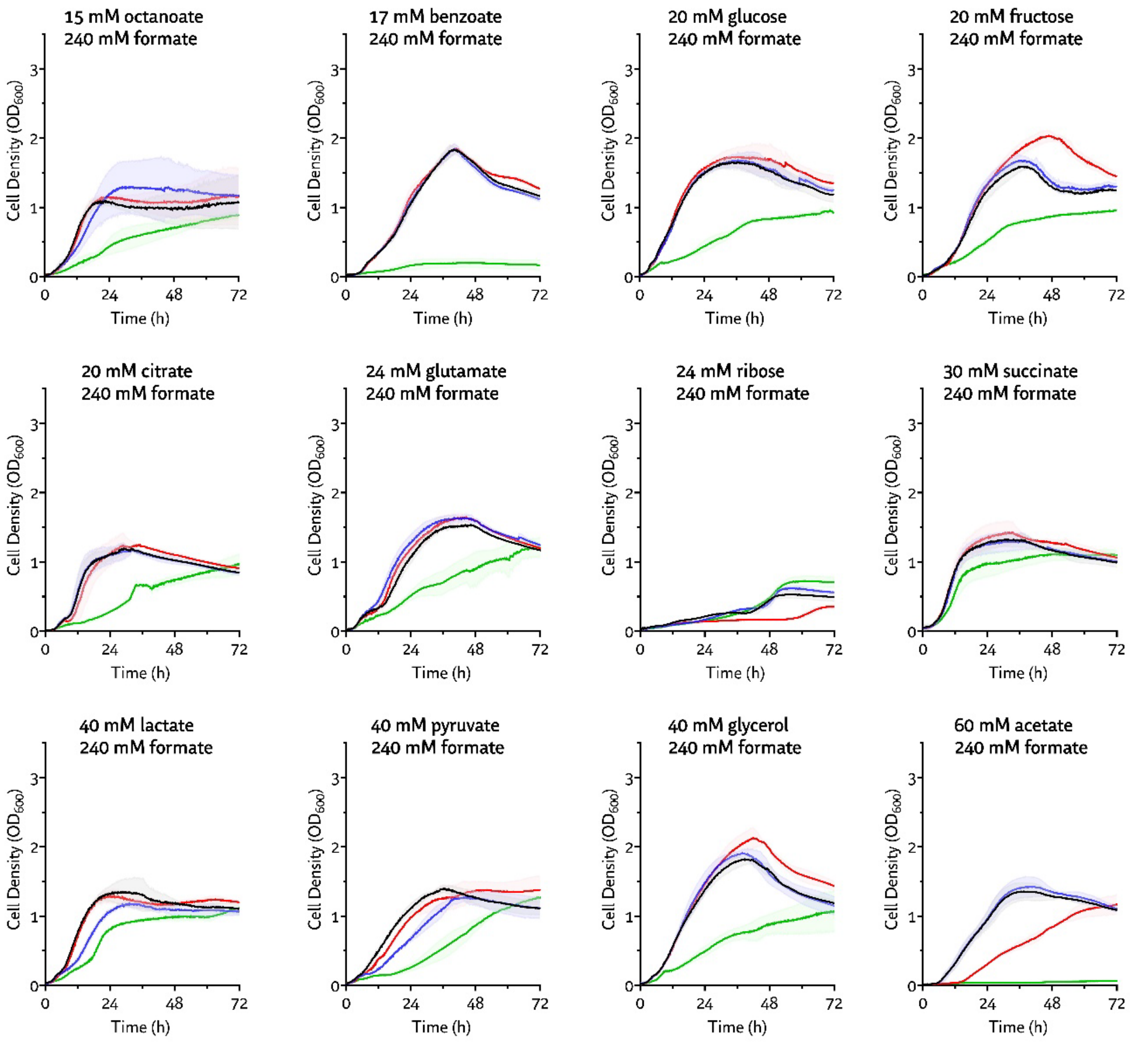


**Figure S6.** Growth curves of *P. putida* KT2440 WT and transhydrogenase mutant strains on different carbon sources supplemented with 240 mM formate. The color code is the same used in the main text (WT in black, Δ*pntAB* in blue, Δ*sthA* in red, Δ*pntAB* Δ*sthA* in green). The experiments were performed in biological quadruplicate (*n* = 4, errors represented as standard deviation).

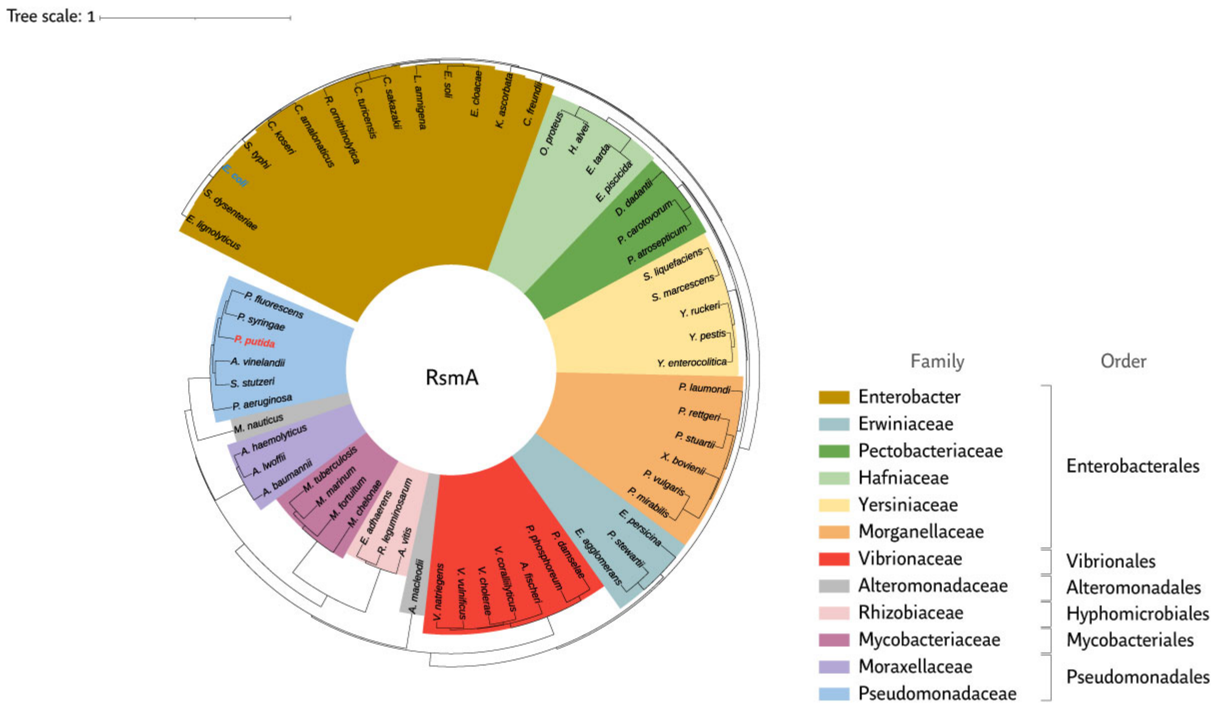


**Figure S7.** Phylogenetic tree of the ribosomal RNA small subunit methyltransferase A RsmA from representative bacteria sharing the dual transhydrogenase adaptation.

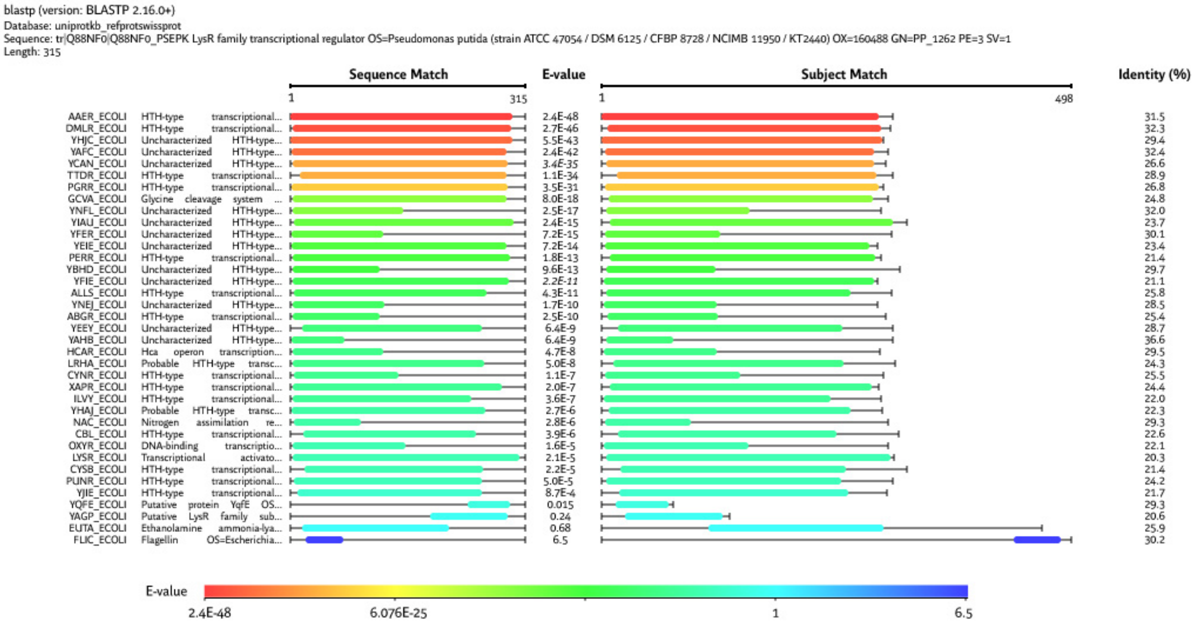


**Figure S8.** Sequence similarity search for PP_1262 in *Escherichia coli* K-12. The sequence of LysR family transcriptional regulator PP_1262 from *P. putida* KT2440 (UniProt entry: Q88NF0) was used as a query for searching the UniProt database via BLAST. The resulting 37 hits are here shown with their names, sequence matches, e-values, subject matches, and percentages of sequence identity.

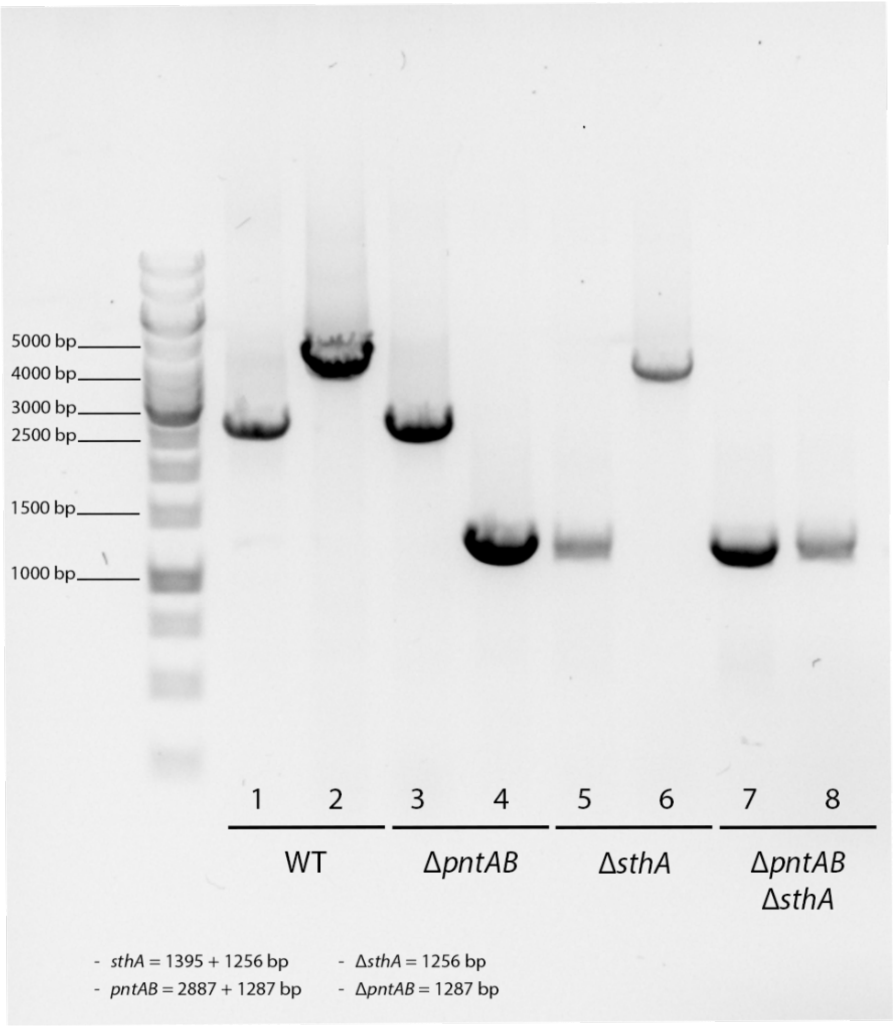


**Figure S9.** Verification of the transhydrogenase-mutant strains Δ*pntAB*, Δ*sthA* and Δ*pntAB* Δ*sthA* by colony PCR. Odd lanes show the amplification of *sthA* with primers 007 and 008 from *P. putida* WT, Δ*sthA*, Δ*pntAB* Δ*sthA* and Δ*pntAB* Δ*sthA* (respectively lanes 1, 3, 5 and 7). Even lanes display the amplification of *pntAB* with primers 009 and 010 from the abovementioned strains (respectively lanes 2, 4, 6 and 8). On the left, the molecular size of the bands of the marker are shown. The expected molecular weights of amplicons with or without deletions are reported in the lower part of the panel.

#### Supplementary References


Bagdasarian, M., Lurz, R., Rückert, B., Franklin, F. C. H., Bagdasarian, M. M., Frey, J., Timmis, K. N., 1981. Specific purpose plasmid cloning vectors. II. Broad host range, high copy number, RSF1010-derived vectors, and a host-vector system for gene cloning in *Pseudomonas*. Gene. 16, 237–247.

Kessler, B., de Lorenzo, V., Timmis, K. N., 1992. A general system to integrate *lacZ* fusions into the chromosomes of Gram-negative eubacteria: regulation of the *Pm* promoter of the TOL plasmid studied with all controlling elements in monocopy. Mol. Gen. Genet. 233, 293–301.

Nikel, P. I., Pérez-Pantoja, D., de Lorenzo, V., 2016. Pyridine nucleotide transhydrogenases enable redox balance of *Pseudomonas putida* during biodegradation of aromatic compounds. Environ. Microbiol. 18, 3565–3582.

Partipilo, M., Ewins, E. J., Frallicciardi, J., Robinson, T., Poolman, B., Slotboom, D. J., 2021. Minimal pathway for the regeneration of redox cofactors. JACS Au. 1, 2280–2293.

Partipilo, M., Whittaker, J. J., Pontillo, N., Coenradij, J., Herrmann, A., Guskov, A., Slotboom, D. J., 2023. Biochemical and structural insight into the chemical resistance and cofactor specificity of the formate dehydrogenase from *Starkeya novella*. FEBS J. 290, 4238–4255.

Platt, R., Drescher, C., Park, S. K., Phillips, G. J., 2000. Genetic system for reversible integration of DNA constructs and *lacZ* gene fusions into the *Escherichia coli* chromosome. Plasmid. 43, 12–23.

Turlin, J., Puiggené, Ò., Donati, S., Wirth, N. T., Nikel, P. I., 2023. Core and auxiliary functions of one-carbon metabolism in *Pseudomonas putida* exposed by a systems-level analysis of transcriptional and physiological responses. mSystems. 8, e00004–00023.

Volke, D. C., Friis, L., Wirth, N. T., Turlin, J., Nikel, P. I., 2020. Synthetic control of plasmid replication enables target- and self-curing of vectors and expedites genome engineering of *Pseudomonas putida*. Metab. Eng. Commun. 10, e00126.

Wirth, N. T., Funk, J., Donati, S., Nikel, P. I., 2023. *QurvE*: user-friendly software for the analysis of biological growth and fluorescence data. Nat. Protoc. 18, 2401–2403.
